# Experimental and Computational Insights into the Structural Dynamics of the Fc Fragment of IgG1 Subtype from Biosimilar VEGF-Trap

**DOI:** 10.1101/2024.08.18.608484

**Authors:** Ebru Destan, Engin Turkut, Alper Aldeniz, Jungmin Kang, Takehiko Tosha, Makina Yabashi, Barış Yılmaz, Ahmet Can Timucin, Hiroaki Matsuura, Yoshiaki Kawano, Irfan Cinkaya, Hasan DeMirci

**Affiliations:** Department of Molecular Biology and Genetics, Koc University, 34450, Istanbul, Türkiye; Biotechnology, DEVA Holding AS, Istanbul, Türkiye; RIKEN SPring-8 Center, Kouto 1-1-1, Sayo-gun, Sayo-cho, Hyogo 679-5148, JAPAN; Graduate School of Science, University of Hyogo, Hyogo, Japan; Japan Synchrotron Radiation Research Institute (JASRI), Kouto 1-1-1, Sayo-gun, Sayo-cho, Hyogo 679-5198, JAPAN; Department of Molecular Biology and Genetics, Acibadem University, Istanbul, Türkiye; Stanford PULSE Institute, SLAC National Laboratory, 94025, Menlo Park, CA, USA; Koc University Isbank Center for Infectious Diseases (KUISCID), Koc University, 34010, Istanbul, Türkiye

## Abstract

The constant fragment (Fc) of the Immunoglobulin G1 (IgG1) subtype is increasingly recognized as a crucial scaffold in the development of advanced therapeutics due to its enhanced specificity, efficacy, and extended half-life. A prime example is VEGF-Trap (Aflibercept), a recombinant fusion protein that merges the Fc region of the IgG1 subtype with the binding domains of vascular endothelial growth factor receptors (VEGFR)-1 and VEGFR-2. The Fc region’s role in N-glycosylation is particularly important, as it significantly influences protein stability. In this study, we present the first near-physiological temperature structures of the N-glycan-bound Fc fragment of IgG1 subtype from a biosimilar VEGF-Trap, determined using the SPring-8 Angstrom Compact free electron LAser (*SACLA*) and the Turkish Light Source (*Turkish DeLight*). Comparative analysis with cryogenic structures, including our own data, reveals alternate conformations within the glycan-binding pocket. Furthermore, molecular dynamics (MD) simulations highlight an unexpected degree of structural plasticity. These findings offer new insights into the molecular basis of Fc-mediated functions and provide valuable information for the design of next-generation therapeutics.

## Introduction

Biology involves a myriad of highly dynamic processes, and a comprehensive understanding of the structural dynamics occurring at the atomic level and near-physiological conditions is a necessity and still remains a significant challenge. X-ray crystallography is a very robust and powerful technique allowing the determination of the arrangement of atoms in a crystal to reveal the three-dimensional structures of biological macromolecules**^1^**. Recent developments of X-ray free electron lasers (XFELs) and sample delivery methods have expanded the boundaries in structural biology**^2,3,4,5^**. These advancements provided new insight into the treatment of diseases in medicine. Here, the information obtained from high-resolution structures of biomolecules can be used in drug design studies, leading to a better understanding of the action mechanism for a potential treatment**^6,7^**.

Since the first discovery of immunoglobulin-like protein by Bence Jones in 1845, the complex nature of immunoglobulins continues to unveil new frontiers in science**^8^**. Immunoglobulins, known as antibodies, are glycoproteins produced primarily by B cells in the adaptive immune system**^9^**. These Y-shaped proteins play a pivotal role in the immune defense mechanism, specifically by recognizing, binding to foreign substances/molecules and neutralizing them or targeting them for destruction. An immunoglobulin involves the attachment of antigen-binding fragment (Fab) and constant fragment (Fc) through the hinge region**^10^**. The polypeptide chains are known as heavy and light chains composed of variable (V) and constant (C) regions**^11^**. While the V region contributes to the antigen-binding site, the C region determines the isotype. The exons encoding the V regions are known as V, D (diversity), or J (joining) genes. The random recombination of the V(D)J gene is the key mechanism to produce antibody diversity**^12^**. Furthermore, somatic hypermutation introduces additional diversity by inducing random mutations in the antibody genes, leading to subtle changes in the antibody’s specificity**^13^**. There are five isotypes of immunoglobulins (Ig): IgA, IgD, IgE, IgG, and IgM. Among them, IgG (specifically the IgG1 subtype) is the most abundant Ig isotype within the blood with its extremely long serum half-life. This makes IgG a suitable scaffold for the design of novel therapeutics**^14,15^**.

Wet Age-Related Macular Degeneration (AMD) is the disease caused by the formation of new anomalous blood vessels driven by angiogenesis in the retina through choroidal neovascularization (CNV)**^16^**. As vascular endothelial growth factor (VEGF), pro-angiogenic cytokines, have a vital role in the progression of angiogenesis, the anti-angiogenic agents that target it have been designed**^17,18^**. VEGF-Trap is a recombinant fusion protein that was first approved by the US Federal Drug Administration (FDA) for the treatment of wet Age-Related Macular Degeneration (AMD) on 18 November 2011**^19,20^**. Its dimeric structure is composed of two chains, the binding domains of vascular endothelial growth factor receptor (VEGFR)-1(Ig2 region) and VEGFR-2(Ig3 region) fused with the Fc fragment of IgG1 subtype **(Supplementary Fig. 1a)**.

The Fc fragment of the IgG1 subtype mediates effector functions of the immune system including Fc receptor binding, antibody-dependent cellular cytotoxicity (ADCC), antibody-dependent cellular phagocytosis (ADCP), and structure stability**^21^**. Compared to all subtypes (IgG1, IgG2, IgG3 and IgG4), IgG1 has the highest affinity to Fcγ receptor (FcγR)**^22^**. Additionally, the Fc fragment contributes to extended drug half-life, improving drug availability and patient compliance. Its capacity for immunomodulation, achieved through interactions with FcγR on immune cells, is particularly valuable in drug development for autoimmune diseases**^23,24^**. Moreover, N-glycosylation in the Fc fragment is a pivotal post-translational modification that significantly influences the diversification of antibody function**^25^**. The addition of glycans to the Fc fragment contributes to the interaction with FcγR**^26,27^**. Mature Fc glycoforms are N-linked and involve seven saccharides: 4 N-acetylglucosamine (GlcNAc) and 3 mannose (Man) residues **(Supplementary Fig. 1b)**. The core glycans can be modified further by the addition of sugars including core fucose (Fuc), bisecting GlcNAc, galactose (Gal) at one or both arms and in the presence of galactose, N-acetylneuraminic acid (NeuAc) or sialic acid**^28^**.

In this study, we revealed four crystal structures of the Fc fragment of IgG1 subtype from biosimilar VEGF-Trap by using three different X-ray sources: Home X-ray source diffractometer (XRD) (Turkish Light Source (*Turkish DeLight*)), Synchrotron (Super Photon ring-8 GeV (*SPring-8*)), and XFEL (SPring-8 Angstrom Compact free electron LAser (*SACLA*)). Rigaku Oxford Diffraction XtaLAB Synergy-S diffractometer equipped with *XtalCheck-S* plate reader system provides rapid and high-quality data collection from protein crystals at ambient temperature. Serially collected diffraction data from low-cost 72-well Terasaki^TM^ crystallization plates using a multiwell-multicrystal plate reader with less amount of crystals offers a cheaper and readily available alternative *in-house* method to serial femtosecond X-ray crystallography (SFX) techniques performed at XFELs**^29^**. The advent of bright X-ray sources such as synchrotron and XFEL facilities has unveiled challenges in protein crystallography, notably pertaining to radiation-induced damage incurred during X-ray irradiation**^30^**. The *SPring-8/SACLA* experimental facility in Japan enables users to perform macromolecular X-ray crystallography experiments with powerful X-rays. The beamline BL32XU with the goniometer-based fully automated data collection system at the *SPring-8* synchrotron radiation facility provides high-throughput diffraction data at cryogenic temperature**^31,32^**. Although cryogenic data collection is a commonly used approach for limiting radiation damage, freezing of a crystal can perturb crystal lattice and protein structures**^33,34^**. A near-physiological crystallographic method that provides high spatial resolution data free of radiation damage and consumes less crystal has been desired. Here, SFX represents an important advancement in protein crystallography as it allows data collection at ambient temperature from a crystal slurry and mitigates radiation damage through the use of ultra-short femtosecond and ultrabright X-ray pulses**^2,35^**. *SACLA* Beamline 2 experimental station has a measurement system with a 10 fs pulse duration and 30 Hz or 60 Hz pulse repetition rate depending on the operation mode, which consists of components including the sample chamber, injector, detector, and data acquisition system**^36^**. For the sample delivery, highly viscous carriers have a great advantage compared to liquid-jet injectors as the sample consumption can be reduced by lowering the flow rate. The high-viscosity cartridge-type (HVC) injector has been designed for SFX experiments equipped in the Diverse Application Platform for Hard X-ray Diffraction in *SACLA* (DAPHNIS)**^4,37^**.

Together with highlighting the structural dynamics of the Fc fragment of the IgG1 subtype, this study provides an invaluable opportunity to compare the abilities of these X-ray sources. In addition to X-ray crystallography experiments, molecular dynamics (MD) simulations were performed to reveal the detailed interaction mechanism of N-glycans with the Fc fragment of the IgG1 subtype. The findings of this study hold the promise of paving the way for designing novel therapeutics with enhanced efficacy.

## Results and discussion

### Near-physiological-temperature structures reveal conformational flexibility occurring through N-glycosylation

We determined two ambient-temperature crystal structures of the Fc fragment of the IgG1 subtype from the biosimilar VEGF-Trap molecule by using two different X-ray sources. Both radiation damage-free XFEL SFX Fc^SFX^ and in-house XRD Fc^XRD(Ambient)^ crystal structures were obtained in orthorhombic crystal form, *P*2_1_2_1_2_1_ space group at 2.00 Å and 2.60 Å resolutions, respectively **(Fig. 1a,b and Supplementary Table 1)**. The well-defined electron density map was observed in the N-glycosylation region where N-glycans bound to Fc fragment of IgG1 subtype: G1F* (chain A) and G0F-GlcNAc (chain B) (GlyTouCan IDs: G56907CZ & G45889JQ) **(Supplementary Fig. 3)**. Moreover, the glycan profile that binds to the T32 site (N_282_) was revealed by using Mass spectrometry (MS) **(Supplementary Fig. 4)**. Based on the MS data, G1F* was observed with higher intensity compared to G0F-GlcNAc. The glycan-binding pocket of two ambient-temperature Fc^SFX^ and Fc^XRD(Ambient)^ structures are analyzed and better electron density was observed for the Fc^SFX^ structure **(Fig. 2a,c)**. For both structures, G1F* glycan was detected more weakly bound to chain A of the Fc fragment compared to GOF-GlcNAc which binds to chain B. B-factor analysis was performed to reveal flexible regions, and chain A is observed as more flexible compared to chain B, leading to the less-defined electron density for G1F* glycan **(Fig. 3)**. The comparison of ambient temperature structures Fc^SFX^ and Fc^XRD(Ambient)^ with previously published Fc^G0&G0F^ structure indicated that loop regions near the glycan-binding site are highly flexible while the core region maintains its stability. Although N-glycosylation is known for increased protein stability, there are cases where N-glycosylation causes flexibility in the structure by inducing altered conformations**^38,39,40^**. Based on the Fc^Deglycosylated^ structure, the binding of glycan causes backbone displacements, inducing changes in protein thermal stability. An important entropic contribution comes from the flexibility of the loop conformation in the glycan-bound state.

**Fig. 1:**
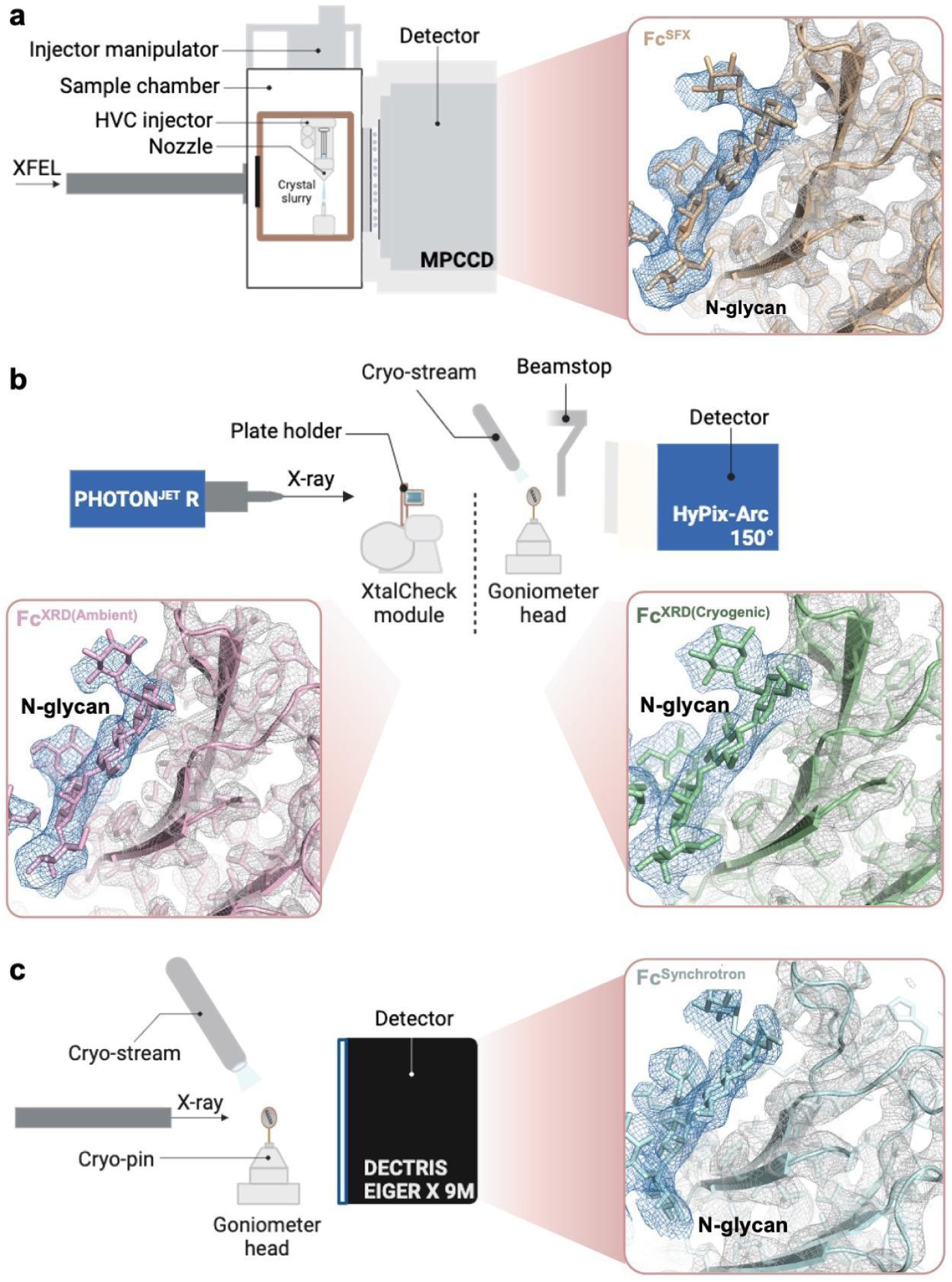
Crystal structures of Fc fragments of IgG1 subtype from biosimilar VEGF-Trap fusion protein. **a** The 2.0 Å crystal structure of the Fc fragment (Fc^SFX^) at ambient temperature is obtained by the SFX technique at SACLA. **b** The 2.6 Å crystal structure of the Fc fragment (Fc^XRD(Ambient)^) at ambient temperature is obtained from *Turkish DeLight* while the 3.14 Å crystal structure (Fc^XRD(Cryogenic)^) was obtained at cryogenic temperature. **c** The 2.64 Å crystal structure of the Fc fragment (Fc^Synchrotron^) at cryogenic temperature is obtained at Spring-8. The *2Fo-Fc* electron densities are contoured at 1 σ level and colored in skyblue (N-glycans) and gray. Fc^SFX^, Fc^XRD(Ambient)^, Fc^Synchrotron^ and Fc^XRD(Cryogenic)^ structures are colored wheat, lightpink, palecyan and palegreen, respectively. PDB IDs and GlyTouCan IDs are indicated in **Supplementary Table 2**.

**Fig. 2:**
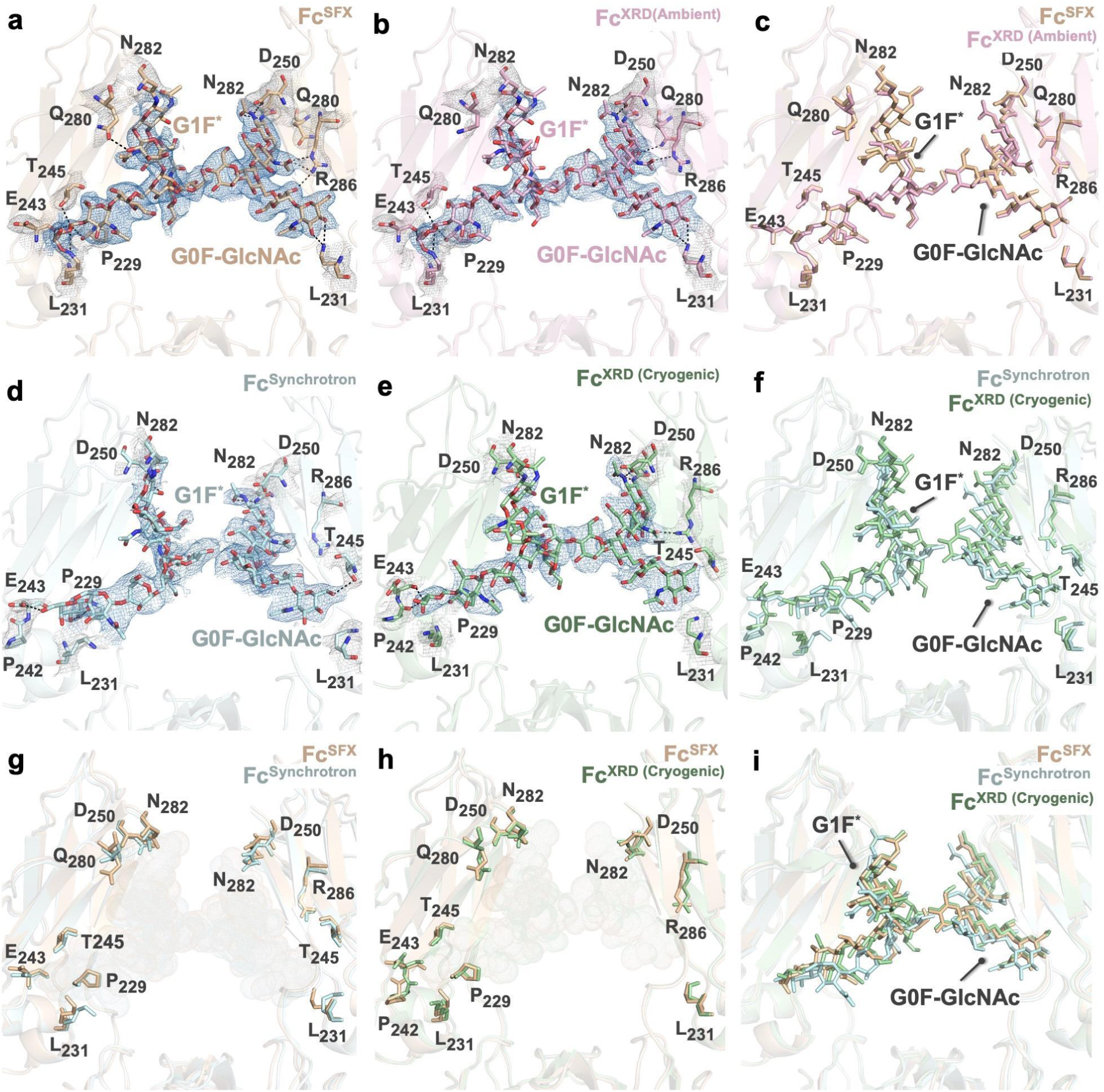
The representation of glycan-binding sites. **a,b** Glycan-binding sites for the Fc^SFX^ and Fc^XRD(Ambient)^ structures are represented to detect the conformational changes between two ambient temperature structures. **c** Fc^SFX^ and Fc^XRD(Ambient)^ structures are superposed with an RMSD value of 0.144 Å. **d,e** Glycan-binding sites for the Fc^Synchrotron^ and Fc^XRD(Cryogenic)^ structures are represented to detect the conformational changes between ambient and cryogenic temperature structures. **f** Fc^SFX^ and Fc^XRD(Ambient)^ structures are superposed with an RMSD value of 0.789 Å. **g-i** Glycan-binding sites for ambient and cryogenic temperature structures are compared to detect the conformational changes. Fc^SFX^ and Fc^Synchrotron^ structures are superposed with an RMSD value of 0.672 Å while Fc^SFX^ and Fc^XRD(Cryogenic)^ structures are superposed with an RMSD value of 0.433 Å. Polar contacts are shown with black dashed lines. The *2Fo-Fc* electron densities are contoured at 1 σ level, and colored in skyblue (N-glycans) and gray (protein residues). Fc^SFX^, Fc^XRD(Ambient)^, Fc^Synchrotron^ and Fc^XRD(Cryogenic)^ structures are colored wheat, lightpink, palecyan and palegreen, respectively. PDB IDs and GlyTouCan IDs are indicated in **Supplementary Table 2**.

**Fig. 3:**
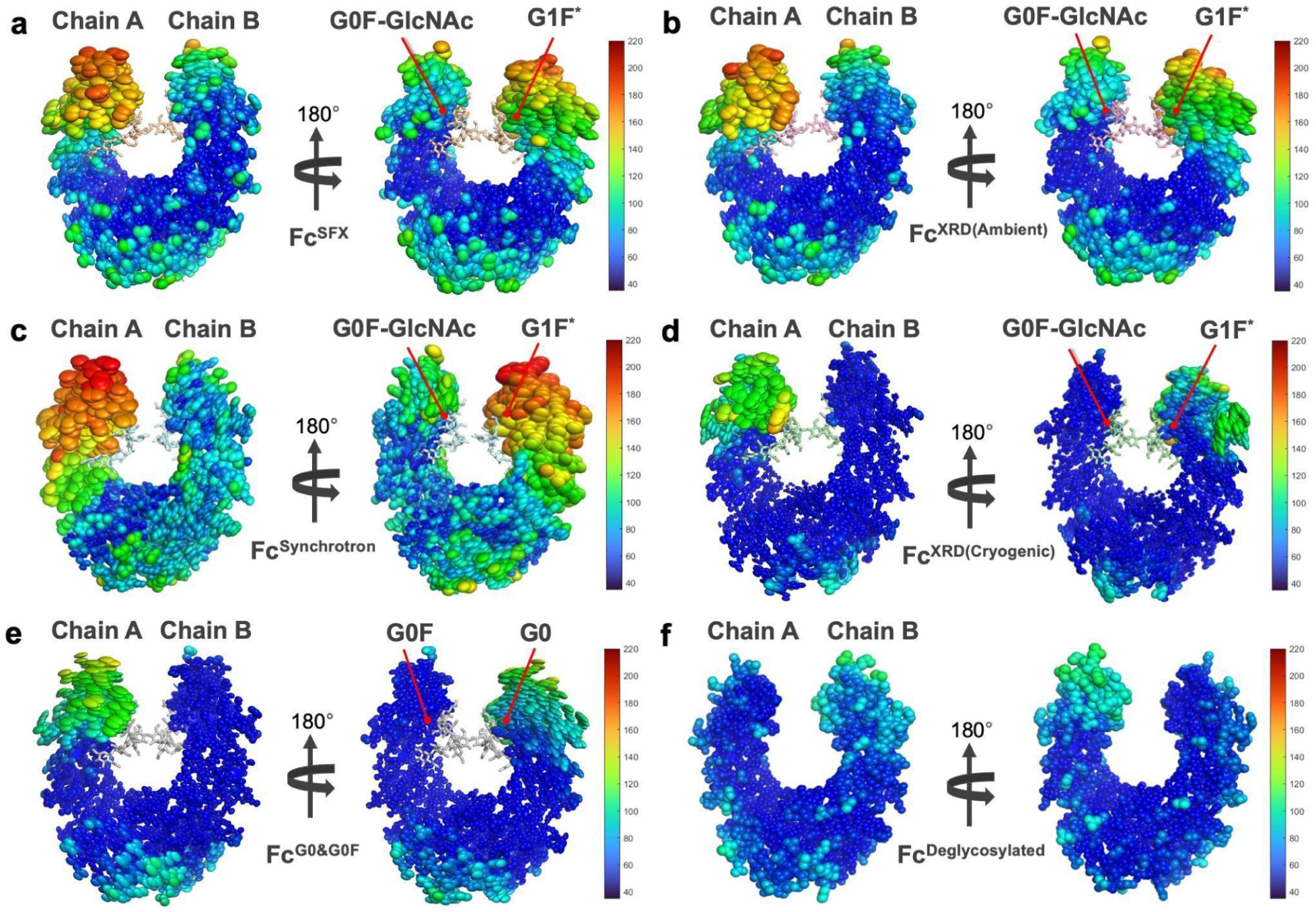
Ellipsoid representation of the crystal structures of the Fc fragment of IgG1 subtype. Each Fc structure is visualized with ellipsoid representation and colored based on b-factors (Spectrum: range (35.00000 to 220.00000)). Ellipsoid representation is used to indicate the flexibility of the macromolecules. RABDAM analysis for Fc^SFX^, Fc^XRD(Ambient)^, Fc^Synchrotron^ and Fc^XRD(Cryogenic)^ structures are summarized in **Supplementary** Fig. 7. PDB IDs and GlyTouCan IDs are indicated in **Supplementary Table 2**.

Two ambient-temperature Fc^SFX^ and Fc^XRD(Ambient)^ structures were superposed with the RMSD value 0.144 Å and only minor conformational changes were observed between the two structures, suggesting that the Rigaku XRD offers high-quality data collection at ambient temperature **(Fig. 2c and Supplementary Fig. 5a,c)**. In the Fc^SFX^ structure, N-glycans interact with N_229_, L_231_, E_243_, T_245_, Q_280_, and N_282_ residues of chain A; and L_231_, D_250_, N_282_, and R_286_ residues of chain B. For the Fc^XRD(Ambient)^ structure, the same polar contacts are observed except the interaction with Q_280_ residue was observed between G0F-GlcNAc rather than G1F*. Furthermore, it is observed that the number of water molecules (Wat) around the glycan-binding site differs between Fc^SFX^ and Fc^XRD(Ambient)^ structures (**Supplementary Fig. 6**). The conserved hydrogen binding network may play a key role during the binding of N-glycans**^41^**. Wat_5_, Wat_111_, and Wat_126_ for Fc^SFX^ structure; Wat_8_, Wat_62_, and Wat_119_ for Fc^XRD(Ambient)^ were observed as conserved and performed hydrogen bond interaction with the critical residues during the interaction with N-glycans. The same conserved water molecule (Wat_702_) was observed for long-chain N-glycan (G1F) in the previously published structure (PDB ID: 5VGP)(Fc^G1F&G0F^). To detect the alternate conformations, the Fc^SFX^ and Fc^XRD^ structures are superposed with previously published structures **(Fig. 4 and Supplementary Fig. 7)**. While only minor conformation changes were observed in the active site residues for the Fc^G0&G0F^ structure, major conformational changes were observed for Fc^G1F*-GlcNAc^ (PDB ID: 1H3V) and Fc^G1F&G0F^ structures (**Fig. 5 and Supplementary Fig. 8a,b)**. Especially, significant conformational changes were observed on L_231_, T_245_, D_250_, Q_280_, N_282_, and R_286_ residues. The comparison of glycan-binding pockets for long and short-chain glycans revealed a significant shift in the position of N-glycans and alternate conformations for the sugar molecules which are the building blocks of N-glycans.

**Fig. 4:**
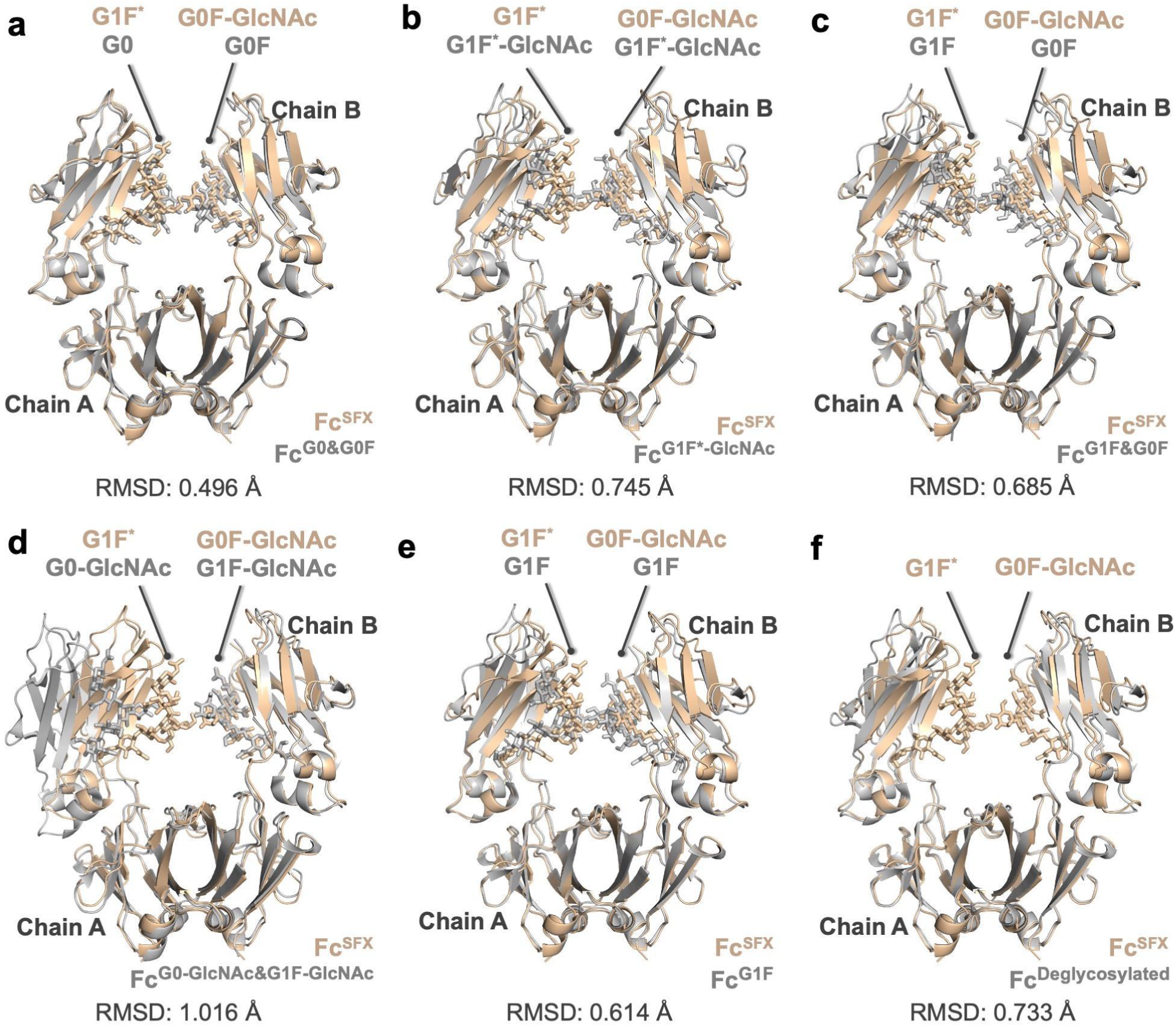
Conformational changes between Fc^SFX^ and cryogenic temperature structures of Fc fragment of IgG1 subtype. Fc^SFX^ structure is superposed with previously published cryogenic temperature structures of the Fc fragment of the IgG1 subtype. Fc^SFX^ structure is colored in wheat while superposed structures are colored in gray. PDB IDs and GlyTouCan IDs are indicated in **Supplementary Table 2**.

**Fig. 5:**
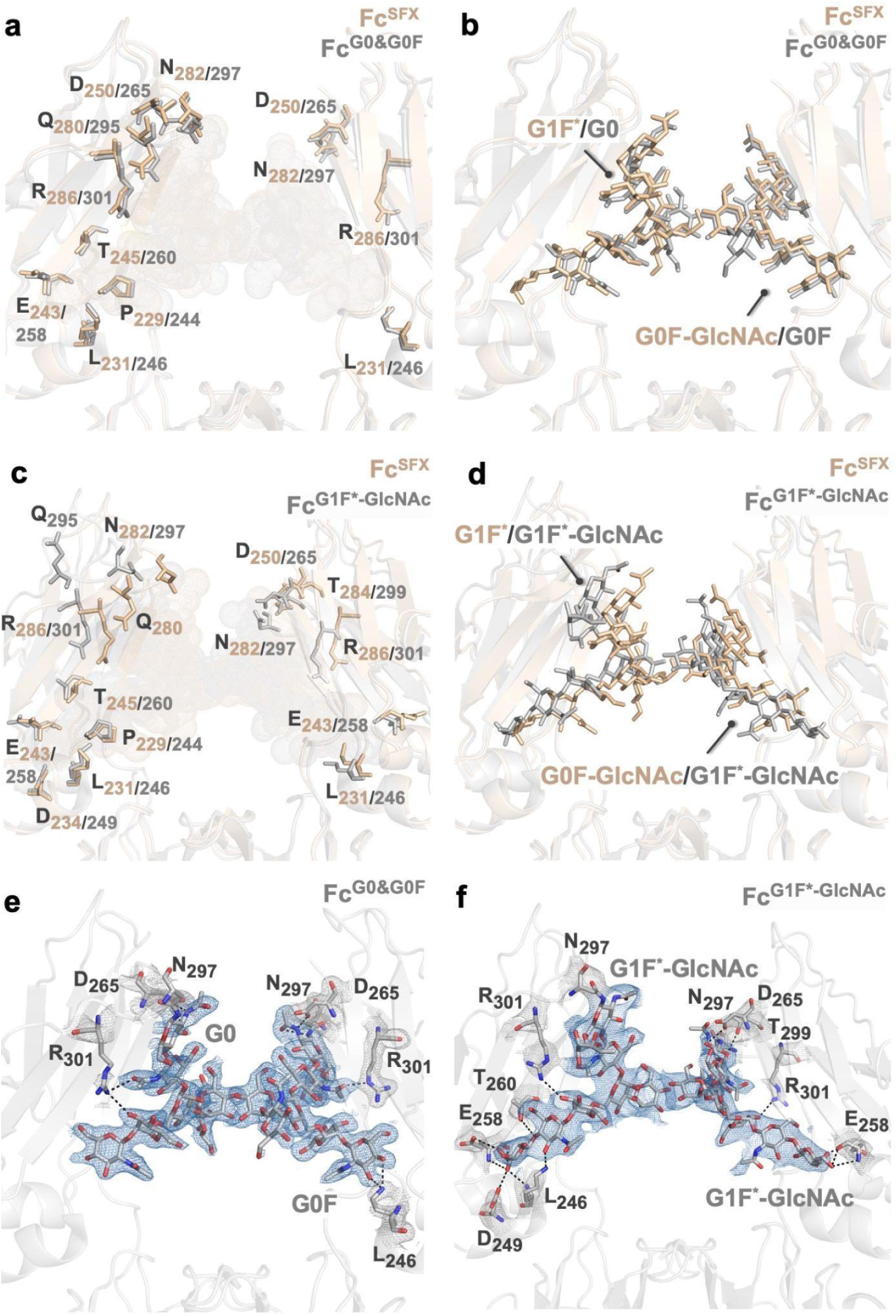
Comparison of the glycan-binding site of Fc^SFX^ with Fc^G0&G0F^ and Fc^G1F*-GlcNAc^ structures (PDB IDs:4W4N&1H3V). Polar contacts are shown with black dashed lines. The *2Fo-Fc* electron densities are contoured at 1 σ level, and colored in skyblue (N-glycans) and gray (protein residues). Fc^SFX^ structure is colored wheat while Fc^G0&G0F^ and Fc^G1F*-GlcNAc^ structures are colored gray. PDB IDs and GlyTouCan IDs are indicated in **Supplementary Table 2**.

### Alternate conformations are observed in the cryogenic-temperature structures

Fc^Synchrotron^ and Fc^XRD(Cryogenic)^ structures were determined at 2.64 and 3.14 Å resolutions from different X-ray sources in orthorhombic crystal form, *P*2_1_2_1_2_1_ space group at cryogenic temperature **(Fig. 1b,c and Supplementary Table 1)**. Here, the difference between the data from multiple crystals and a single crystal impacted the overall structural dynamics. Commonly, both cryogenic temperature structures displayed significant conformational changes in the glycan-binding site, highlighting the importance of structural dynamics at near-physiological temperatures **(Fig. 2 and Supplementary Fig. 9)**. Previously observed common hydrogen bond interactions between the N-glycans and L_231_ (chain A&B), T_245_ (chain A), D_250_ (chain B), and R_286_ (chain B) residues were not observed for the Fc^Synchrotron^ structure. The major conformational changes were observed on the L_231_ and Q_280_ (weak electron density for Fc^Synchrotron^ structure) residues in the chain A while L_231_, D_250,_ and N_282_ in the chain B **(Fig. 2g)**. For the Fc^XRD(Cryogenic)^ structure, G1F* interacts with N_229_, P_242_, E_243_, D_250_, and N_282_ residues of chain A while G0F-GlcNAc interacts with D_250_, R_286,_ and N_282_ residues of chain B. The major conformational changes were observed on the L_231_, P_242_, E_243_ and Q_280_ (weak electron density for Fc^XRD(Cryogenic)^ structure) residues in chain A while L_231_ and R_286_ in chain B **(Fig. 2h)**. Interestingly, the comparison of glycan-binding positions indicated the altered N-glycan confirmation for cryogenic structures, especially Fc^Synchrotron^ structure **(Fig. 2i)**. To highlight the structural dynamics and elucidate the radiation damage from different X-ray sources, b-factors were analysed by using *RABDAM* software**^42^ (Fig. 3 and Supplementary Fig. 10)**. *RABDAM* generates the B_Damage_ metric using atomic isotropic B-factor values in environments with similar packing density, and the B_net_ metric, which is a derivative of B_Damage_ that summarizes the extent of specific radiation damage. Accordingly, both cryogenic and ambient temperature structures have low radiation damage values (B_net_= 0.9-1.5) based on data collection set up**^43^**. These findings indicate the high b-factor values mostly correlated with the mobility of the structure.

Fc^Synchrotron^ and Fc^XRD(Cryogenic)^ structures are superposed with previously published cryogenic temperature structures **(Supplementary Fig. 11,12)**. Based on the comparison of Fc^Synchrotron^ and Fc^G0&G0F^ structures, minor conformational changes with 0.604 Å RMSD were observed in the glycan-binding site residues, except for the L_231_ residue **(Supplementary Figure 13c)**. Apart from Fc^G0&G0F^structure, major conformational changes up to 0.922 Å were observed in Fc^G1F*-GlcNAc^ and Fc^G1F&G0F^ structures **(Supplementary Figs. 8c,14c)**. For the Fc^XRD(Cryogenic)^ structure, similarly minor conformational changes with 0.422 Å RMSD were observed compared to the Fc^G0&G0F^ structure while major conformational changes up to 0.866 Å were observed in Fc^G1F*-GlcNAc^ and Fc^G1F&G0F^ structures **(Supplementary Figure 8e,13e,14e)**. These findings indicate the different structural conformation based on N-glycan composition (Supplementary Fig. 8,13,14).

### Superposition with previously published structures reveals the conformational changes upon binding to FcRn, FcγR, or C1q proteins

The Fc fragment of the IgG1 subtype is a common scaffold for the designing of recombinant proteins. It has binding sites for neonatal Fc receptor (FcRn), FcγR and complement protein C1q for the functioning of the immune system **(Fig. 6a)**. Here, we observed the conformational changes on the interaction site in the absence of FcRn, FcγR, and C1q proteins. Each IgG1 molecule has one binding site for FcγR and two putative binding sites for C1q and FcRn **(Fig. 6b)**. The superposition of our structures with FcRn bound structure (PDB ID: 1FCC) (Fc^FcRn^) highlighted the major conformational changes on the key residues (L_236_, M_237_, I_238_, Q_296_, E_367_ and H_418_) during the binding of FcRn **(Fig. 6c-e; Supplementary Movie 1,2; Supplementary Figs. 15a,d, 16a,d and 17a,d)**. For the FcγR binding site, significant conformational changes were observed in the loop region of chain A near the glycan-binding site **(Fig. 6f-h; Supplementary Movie 3,4; Supplementary Figs. 15b,e, 16b,e and 17b,e)**. The S_224_, D_250_, S_252_, E_254_, and Y_281_ residues in the loop are shifted in the presence of FcγR binding (PDB ID: 7URU) (Fc^FcγR^). Although the C1q protein has two binding sites for chains A and B, the chain A binding site is not visible as it is located in the hinge region. CryoEM structure of C1q-Fc complex structure (PDB ID: 6FCZ) (Fc^C1q^) was superposed with our structures **(Fig. 6i-k; Supplementary Movie 5,6; Supplementary Figs. 15c,f, 16c,f and 17c,f)** and the major conformational changes on the E_254_, H_253_, and S_283_ residues were observed when it is in complex with C1q.

**Fig. 6:**
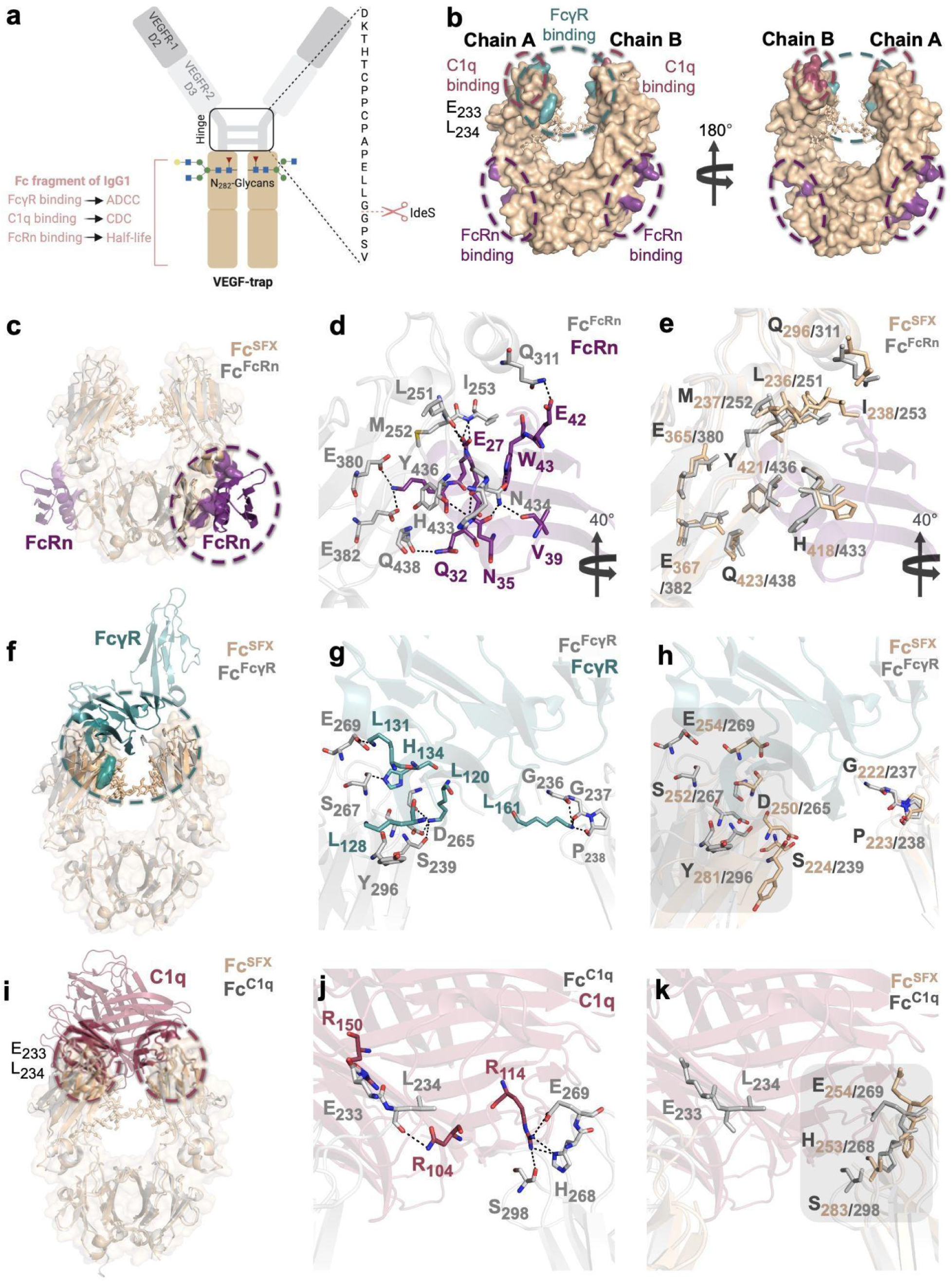
Conformational changes of the active site residues on Fc fragment in the presence of FcRn, FcγR and C1q. **a** The systematic representation of the VEGF-Trap is illustrated to indicate the cleavage site of IdeS protease for obtaining the Fc fragment. **b** Fc fragment involves binding sites for FcRn, FcγR and C1q. Fc^SFX^ structure is colored in wheat while FcRn, Fcγ and C1q binding sites are colored in violetpurple, lightteal and raspberry, respectively. **c-e** Fc fragment of IgG1 subtype in complex with FcRn (Protein G bound) (PDB ID: 1FCC) (Fc^FcRn^) and Fc^SFX^ structures are superposed with RMSD value 0.873 Å. Fc^FcRn^ is colored in gray while FcRn binding region and Protein G are colored in violetpurple. **f-h** Fc fragment of IgG1 subtype in complex with FcγR (PDB ID: 7URU) (Fc^FcγR^) and Fc^SFX^ structures are superposed with RMSD value 0.627 Å. Fc^FcγR^ is colored in gray while FcγR binding region is colored in lightteal. **i-k** Fc fragment of IgG1 subtype in complex with C1q (PDB ID: 6FCZ) (Fc^C1q^) and Fc^SFX^ structures are superposed with RMSD value 1.067 Å. Fc^C1q^ is colored in gray while C1q binding region is colored raspberry. There is one available binding site for FcγR and two potential binding sites for C1q and FcRn per IgG1 molecule. C1q binding site in Chain A is not visible for Fc^SFX^ structure as E_233_ and L_234_ residues are located in the hinge region. Fc^SFX^ structure is colored in wheat. Polar contacts are shown with dashed black lines. As a result of superposition, the major shifts in the secondary structures are indicated with gray squares.

### MD simulations highlight the details of glycan-binding pocket dynamics and inter-domain motions

The comparison of G1F* and G0F-GlcNAc bound structures with the Fc^Deglycosylated^ structure indicated the significant conformational changes in the glycan-binding site (**Supplementary Fig. 18)**. Especially, N_282_ which plays a key role during the N-glycosylation process was detected as a major conformational change, indicating the movement of the loop region during the glycan binding (**Supplementary Movie 7-10**). To highlight the binding mechanism of N-glycans, MD simulations were performed using Fc^SFX^, Fc^Synchrotron,^ and Fc^Deglycosylated^ structures. As we observed altered N-glycan conformation at cryogenic temperature, computational observation of Fc^Synchrotron^ structure offers a new insight due to the effect that the initial structure lacks some glycan-amino acid linkages. RMSD (Å) calculation based on the backbone atoms of each protein indicated the stable maintenance of the conformations **(Fig. 7a and Supplementary Fig. 19a-c)**. To detect the structural changes through 250 ns, a radius of gyration (Å) calculation based on the center of mass of each atom in the molecule was performed, providing initial information about the molecule’s overall shape and compactness. While there is no major difference between Fc^SFX^ and Fc^Deglycosylated^ structures, a small jump was detected in the fiftieth nanosecond for the Fc^Synchrotron^ structure **(Fig. 7b and Supplementary Fig. 19d-f)**. Following the jump, this change in compactness converges back to the initial state, indicating secondary structure stabilization. As the next step, highly flexible regions were revealed by RMSF-CA (Å) calculation. It is observed that three loop regions (I, III, IV) near to glycan-binding site are more flexible for both chains compared to the rest of the molecule **(Fig. 7c,d)**. Within these regions, I (Residue range: 244-262) and IV (Residue range: 306-321) regions in chain A were detected as ∼1.5-fold more flexible compared to the Fc^Deglycosylated^ structure, indicating the effect of long-chain N-glycan (G1F*). For chain B where the short-chain glycan (G0F-GlcNAc) is bound, the loop region II (Residue range: 264-272) was detected as more flexible compared to both the Fc^Deglycosylated^ structure and chain A of Fc^SFX^, Fc^Synchrotron^ structures. These findings indicate that the binding of N-glycans causes additional flexibility on the protein, supporting the previous results from X-ray crystallography data **(Fig. 3)**.

**Fig. 7:**
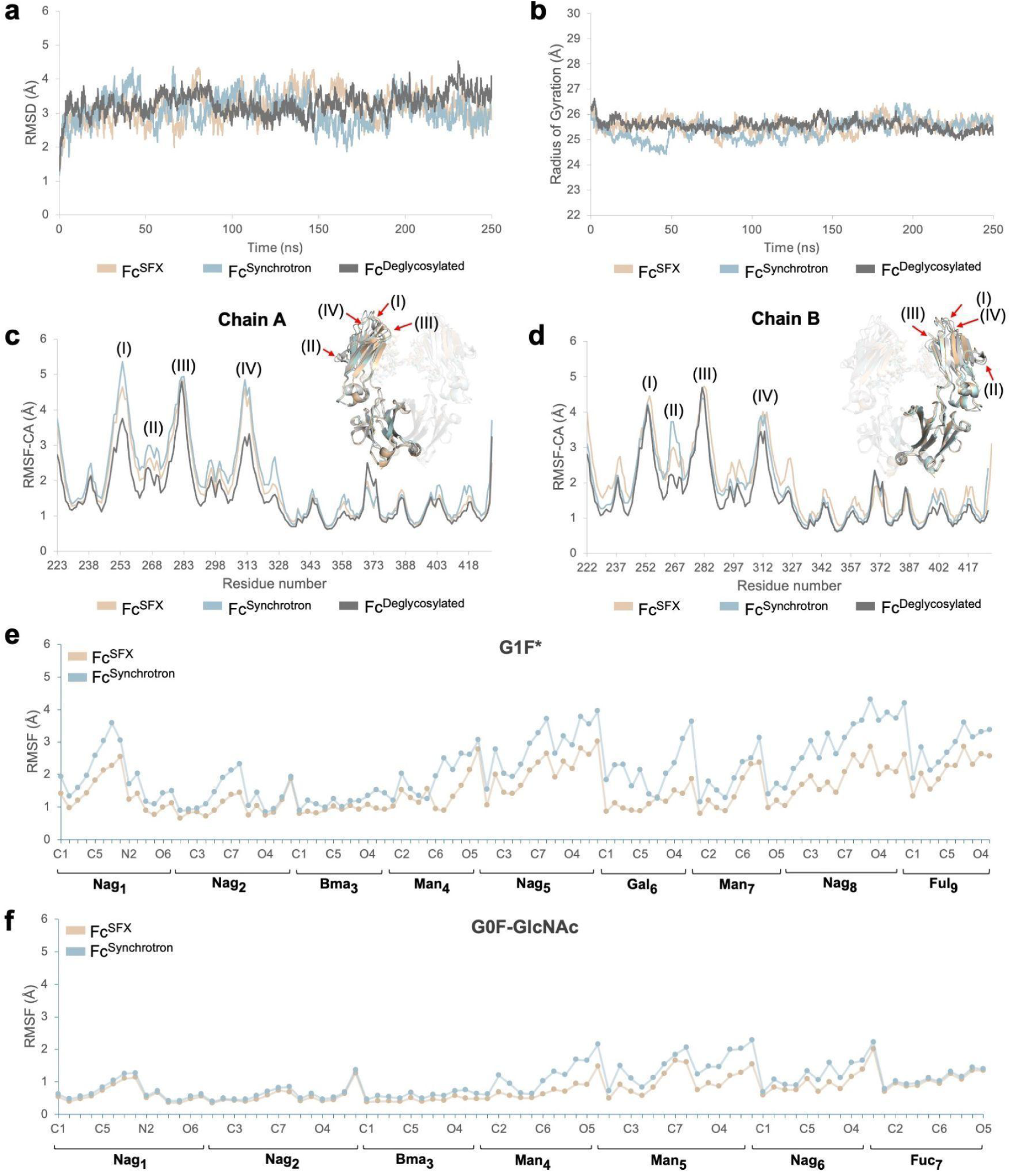
Structural dynamic differences in the presence of N-glycans based on Molecular Dynamic simulations. The result of three production runs (total of 750 ns) is averaged and the deglycosylated Fc fragment from IgG1 subtype (PDB ID: 7RHO)(Fc^Deglycosylated^) is used as a reference structure for the analysis. **a,b** RMSD (Å) and radius of gyration (Å) are calculated for each structure to reveal the protein compactness through the simulation period (250 ns). **c,d** RMSF (Å) is calculated for each chain based on alpha carbons. High-fluctuated regions are indicated with red arrows. **e,f** RMSF (Å) is calculated for N-glycans which are bound to chain A (G1F*) and chain B (G0F-GlcNAc). The calculation is performed for each atom that builds the residues of N-glycan.

Atom versus RMSF (Å) values were calculated for N-glycans to detect fluctuations of atoms during the simulation time frame. **(Fig. 7e-f)**. It is observed that G0F-GlcNAc which bound the chain B is more stable compared to G1F*, explaining the weak electron density for G1F* in the chain A. To provide a better understanding of conformational changes in the presence of N-glycans, the frames of production run 3 (R3) were analyzed in detail **(Supplementary Movie 11-13)**. It is observed that the addition of G1F* causes a significant displacement in chain A (especially the C_H_2 domain) while chain B maintains its stability **(Supplementary Fig. 20,21)**. These dynamics in the presence of N-glycans may induce the binding of other partners to the Fc region in the immune system**^44^**. Based on atom versus RMSF (Å) values, glycan-binding to the critical residues does not interfere with the general trend that surface exposed residues becoming more flexible than buried ones compared with the Fc^Deglycosylated^ structure **(Supplementary Fig. 22a-h)**. However, RMSF-CA (Å) and RMSF-Side Chain (Å) calculations revealed differences, supporting the chain movement in the C_H_2 domain. It is observed that D_250_ is more flexible while N_282_ is more stable in the presence of N-glycans **(Supplementary Fig. 22i-l and Supplementary Table 3)**. Additionally, distances were calculated between the critical residues and N-glycans to determine the stability of the polar contacts **(Supplementary Fig. 23)**. Comparison of distance measurements for the residues P_229_, L_231_, E_243_ and T_245_ in chain A indicated that the flexible G1F* glycan returns back to close position with these residues for Fc^Synchrotron^ structure while it tends to move away for Fc^SFX^ structure. Overall, these counter motions once again show us interaction patterns of flexible G1F\***(Supplementary Fig. 24)**. For chain B, as G0F-GlcNAc is less flexible in the glycan-binding pocket, the distances were detected as stable throughout the simulation period **(Supplementary Fig. 25)**. These findings indicated the unstable state of the long-chain ligand.

## Conclusion

The data presented herein offers a comprehensive analysis of the binding mechanism of N-glycans and its effects on protein structure. Insights into how N-glycans induce conformational changes may enable the engineering of drugs with optimized potency, stability, efficacy, and pharmacokinetics. For this purpose, the near-physiological structure is essential as it closely mimics the near-natural conformation. The experimental and computational observation of the structural dynamics revealed that the binding of long and short-chain N-glycans differently affects the stability of the Fc fragment of the IgG1 subtype from biosimilar VEGF-Trap.

## Methods

### Protein Sample Preparation

Full-length biosimilar VEGF-Trap protein (40 mg/ml) which is produced by our DEVA Holding A.S team in Türkiye was incubated with 10 μL IdeS protease (PROMEGA, USA) for 3 hours at 37 °C to cleave the Fc region below hinge region, generating the binding domains of VEGFR-1 and VEGFR-2 and Fc fragment. Further, the size exclusion chromatography method was applied to purify the ∼50 kDa Fc fragment. Superdex^TM^ 200 (S200) column was used with the buffer containing 150 mM NaCl and 20 mM Tris-HCI (pH 6.2). Elutions were collected for each peak and the Fc fragment peak was inspected using SDS-PAGE. Finally, the pure protein was pooled and concentrated by Amicon ultrafiltration 30 kDa cutoff spin filters from Millipore to a final concentration of 10 mg/ml and stored at -80°C until crystallization trials.

### Crystallization

Sitting-drop vapor diffusion micro-batch under the oil technique was used for initial crystallization screening as explained by Ertem *et al*.**^45^**. Purified Fc protein at 10 mg/ml mixed with approximately 3500 commercially available sparse matrix crystallization screening conditions (1:1 v/v) in a 72-well Terasaki^TM^ plate (Greiner-Bio, Germany). Then, each well was covered with 16.6 µL paraffin oil (Tekkim Kimya, Turkiye) and incubated at room temperature. The best crystals were obtained at Index™ crystallization screen 1 condition #43, containing 0.1 M Bis-Tris (pH 6.5), 25% w/v Polyethylene glycol (PEG) 3350, from Hampton Research, USA **(Supplementary Fig. 2a)**. Replica plates were done by using the same crystallization screen condition. After multiple optimizations of the seeding protocol by using crystals obtained by micro-batch under oil, we scaled up the batch crystallization volume to a total of 1.5 ml. During the scale-up process, the formation of microcrystals was observed by a compound light microscope **(Supplementary Fig. 2b)**.

### Ambient & Cryogenic-temperature data collection and data processing at *Turkish DeLight*

Initial ambient-temperature data was collected by using Rigaku’s XtaLAB Synergy Flow XRD system controlled by *CrysAlisPro* software (Rigaku Oxford Diffraction, 2022) as described by Gul et al.**^29^**. The airflow temperature of Oxford Cryosystems’s Cryostream 800 Plus was adjusted to 300 K (26.85 °C) and the 72-well Terasaki plate was placed on the modified adapter of the *XtalCheck-S* plate reader. After selecting well-diffracting crystals, diffraction data were collected from multiple crystals by using *automatic data collection* mode. During the data collection, the detector distance was set at 100.00 mm and exposure time was 5.00 seconds per image. The diffraction data was collected with different scan widths (0.10, 0.20, and 0.50 degrees) and X-ray intensities (2, 4 and 10%). A total of 14898 frames were collected in 20 hours. The collected data was processed using the *automatic data reduction* process and merged using the *proffit merge* process of *CrysAlisPro* software (Rigaku Oxford Diffraction, 2022). For cryogenic temperature data collection from a single crystal, Oxford Cryosystems’s Cryostream 800 Plus was adjusted to 100 K and data was collected described as Atalay *et al*.**^46^**. A total of 720 frames were collected in 32 min.

### Cryogenic-temperature data collection and processing at *Spring-8*

Diffraction data from multiple crystals was collected at the BL32XU beamline at SPring-8 using the automated data-collection system *ZOO* described by Hirata *et al*.**^32^**. Before the data collection, crystals were frozen in liquid nitrogen and pucks were replaced into the dewar which is filled with liquid nitrogen in the hutch. All data sets were acquired using a continuous helical scheme with 360° of oscillation and the following experimental parameters: oscillation width, 0.1°; exposure time, 0.02 s; beam size, 10 µm (horizontal) × 15 µm (vertical); wavelength, 1 Å; average dose per crystal volume, 10 MGy; detector, EIGER X 9M (Dectris); temperature, 100 K. The data obtained were automatically processed by XDS**^47^** in KAMO**^31^**.

### SFX data collection and processing at *SACLA*

Diffraction data from microcrystals was collected at the Experimental Hutch 3 of *SACLA* Beamline 2 using 10 keV X-rays with a pulse duration of <10 fs and a repetition rate of 30 Hz**^48,49^**. To obtain 10^8^ crystal density for SFX experiments, 500 μL microcrystal slurry was centrifuged at 3000 rpm for 5 min, and 490 μL supernatant was discarded for each injection. The remaining 10 μL crystal slurry with 10^8^ crystal density was mixed with grease (No. 42150, Synco Chemical Co.) on a plastic plate according to a previously reported procedure **(Supplementary Fig. 2c)^50^**. The sample was directly transferred into a 200 μL sample cartridge carefully by using a spatula without leaving any bubbles inside the sample-grease mix. Then, the cartridge was loaded into the HVC injector with a nozzle of 75 μm ID. The HVC injector was installed in a He-gas-filled diffraction chamber of the DAPHNIS instrument**^37^**. Diffraction patterns were recorded using the MPCCD detector at a sample-to-detector distance of 50 mm**^51^**. HPLC flow rate was set as 0.0127 µL min^−1^ and the sample slurry (mixed with grease) was injected at a flow rate of 0.79 µL min^−1^. Diffraction images were filtered to extract ‘hit’ images through the *CHEETAH***^52^** pipeline at *SACLA* and indexed by using *CrystFEL***^53,54,55^**. During the beamtime, 3 injections were done and a total of 24873 out of 66849 hits were indexed. Final data processing and merging were performed using *CrystFEL*.

### Structure determination and refinement

Four structures of the Fc fragment of IgG1 subtype from VEGF-Trap were determined in space group *P*2_1_2_1_2_1_ by using the automated molecular replacement program *PHASER***^56^** implemented in *PHENIX***^57^** suite. The coordinates of previously published cryogenic-temperature structure (PDB ID: 4W4N) (Fc^G0&G0F^) were used for initial rigid body refinement. After simulated-annealing refinement, individual coordinates and TLS parameters were refined. N-glycans are replaced into the structure and altered side chains and water molecules were checked in *COOT***^58^**. The positions with strong difference density were retained and water molecules located outside of significant electron density were manually removed. Data collection and structure refinement statistics are summarized in **Supplementary Table 1**. For the structural alignments and figure generation, *PyMOL***^59^** was used and secondary structure alignment was performed by using *JALVIEW***^60^**. Quantification of radiation damage for each structure was performed by using *RABDAM* software**^42^** and all kernel density plots were drawn by using *MATLAB***^61^**.

### MD simulations and data analysis

All MD simulations were performed by using the *Nanoscale Molecular Dynamic (NAMD 3.0)* program**^62^** with the *CHARMM36m***^63^** force field. Pre-processing of the structures (protein and N-glycans) including the addition of solvation box and ions was completed through *CHARMM-GUI***^64^** Input Generator/Solution Builder. The *TIP3P* model was used for water molecules within the system during the simulation. The system size (cubic) was determined to have at least 10 Å from protein in all axes and 150 mM NaCl was added for neutralization of charge. Protonation states of the histidines were determined using *PDB2PQR* software**^65^**. Disulphide bridges were formed based on the structural information provided within the PDB files. In the deglycosylated cryogenic temperature structure (PDB ID: 7RHO) (Fc^Deglycosylated^) only, protein chains were completed for 2-3 missing amino acid residues in N and C terminal ends, respectively. Systems were fully energy minimized 10000 steps with conjugate gradient method, which was followed by equilibration with 125000 steps of canonical (NVT) ensemble**^66^**. During equilibration, protein positional restraints and carbohydrate dihedral restraints were employed as provided by *CHARMM-GUI* output. For production simulations, an isothermal-isobaric (NPT) ensemble was employed with periodic boundary conditions and coulomb interactions were calculated using the particle-mesh *Ewald* algorithm**^67^**. 10 Å of switching distance and 12 Å of cutoff distance limits were used to calculate non-bonded electrostatic interactions and *Van der Waals* interactions. The pressure was held at 1 atm using the *Nose-Hoover Langevin* method and temperature was kept at 310 K with Langevin dynamics**^68^**. Lengths of the hydrogen-containing bonds were constrained using the *SHAKE* algorithm**^69^**. A 2 fs time step was used in all simulations. The production runs were set as 250 ns in triplicates. In total, 750 ns MD simulation was performed for each structure and the total run time was 2.25 µs.

*VMD* software**^70^** was used for the analysis of trajectories and the visualization of structures. Root mean square displacement (RMSD) calculations were performed to determine conformational stability through 250 ns MD simulations. Before each calculation, the structure was aligned to the initial frame and RMSD was calculated based on the backbone atoms for each production run by using the RMSD trajectory tool module of *VMD*. The radius of gyration was calculated based on the center of mass to characterize the spatial conformation and measure the compactness. The root-mean-square-fluctuation (RMSF) and distance calculations were performed to reveal the details of structural dynamics in the glycan-binding site. All the calculations in the study were computed with tcl scripts. As there are three production runs for each structure, the results were averaged and considered for the interpretation of the data. All simulation movies were generated by using the movie maker module of *VMD*.

## Data availability

Coordinates and structure factors have been deposited in the RCSB Protein Data Bank under accession codes 8ZCK and 8ZCL, which correspond to the ambient temperature structures (Fc^SFX^ and Fc^XRD(Ambient)^, respectively); and 8ZCM and 9IIE, which correspond to the cryogenic structures (Fc^Synchrotron^ and Fc^XRD(Cryogenic)^, respectively). Any remaining information can be obtained from the corresponding author upon request.

## Acknowledgments

This publication has been produced benefiting from the 2244 Industrial PhD Fellowship Program Funding Program of the Scientific and Technological Research Council of Turkey (TÜBİTAK) (Project Numbers: 119C132). However, the entire responsibility of the publication belongs to the owner of the publication. The financial support received from TÜBİTAK does not mean that the content of the publication is approved in a scientific sense by TÜBİTAK. Portions of this research were carried out at the *SPring-8* Angstrom Compact free electron LAser, Japan (*SACLA*) and supported by the *SACLA* Research Support Program for Graduate Students (Proposal numbers: 2023B8058 and 2023B2761). The authors gratefully acknowledge the use of the services and Turkish Light Source (*Turkish DeLight*) X-ray facility at the University of Health Sciences, Experimental Medicine Application & Research Center (DETUAM), Validebag Research Park. We thank Jerome Johnson for his help during cryogenic temperature data collection at *Turkish DeLight*. The numerical calculations reported in this work were fully performed at TUBITAK ULAKBIM, High Performance and Grid Computing Center (TRUBA resources).

## Author Contributions

The project was initiated and coordinated by H.D., I.C.; Fc fragment of VEGF-Trap is purified by E.D., E.T., A.A., B.Y. and I.C.; Diffracting crystals were obtained and optimized by E.D.; Crystallography applications at *SPring-8/SACLA* were supported by E.D., J.K., T.T. and M.Y; and in beamtime, samples were prepared by E.D.; SFX data at *SACLA* was collected by E.D. and J.K.; Synchrotron data at *SPring-8* was collected by E.D., J.K. and T.T with the help of Y.K.; Crystallography applications at *Turkish DeLight* were supported by E.D. and H.D.; and in beamtime, samples were prepared by E.D. and the data collected by E.D. and H.D.; The data processing was performed for SFX and XRD structures by E.D. and for synchrotron structure by H.M.; Structures were refined by E.D.; and data were interpreted by E.D. with the help of H.D. and I.C.; Molecular dynamics simulations were performed by A.C.T.; and data analyzed and interpreted by E.D. and A.C.T. All of the authors read and acknowledged the manuscript.

## Supplementary Information

Supplementary Fig. 1-25; Supplementary Table 1-3; Supplementary Movie 1-13

