## Supplementary material for "Experimental and Computational Insights into the Structural Dynamics of the Fc Fragment of IgG1 Subtype from Biosimilar VEGF-Trap": Supplementary Information.pdf

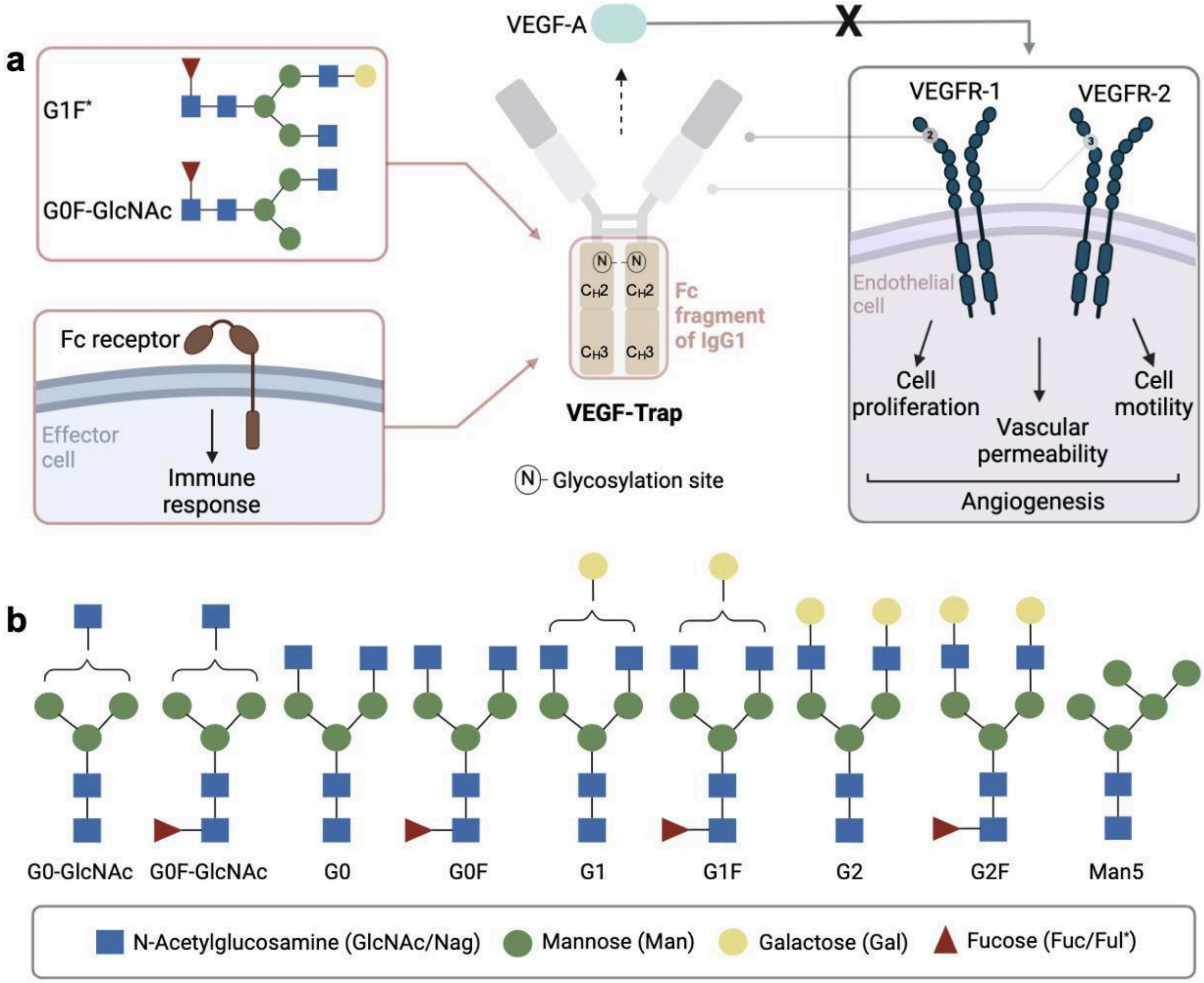


**Supplementary Fig. 1: Fc fragment of IgG1 subtype from biosimilar VEGF-Trap fusion protein. a** The action mechanism of the VEGF-Trap and the role of the Fc fragment for immune response is illustrated. VEGF-Trap consists of the binding domains of VEGFR-1 and VEGFR-2 fused with the Fc fragment of IgG1 subtype (including constant ($C_H2$ and $C_H3$) domains). It targets VEGF-A to block the interaction with its receptor, inhibiting angiogenesis. Fc fragment involves two N-glycosylation sites and a binding site for the Fc receptor, which is colored in wheat. **b** Common N-glycans that bind the Fc fragment of IgG1 subtype are illustrated. This figure is created by using BioRender (https://www.biorender.com/).

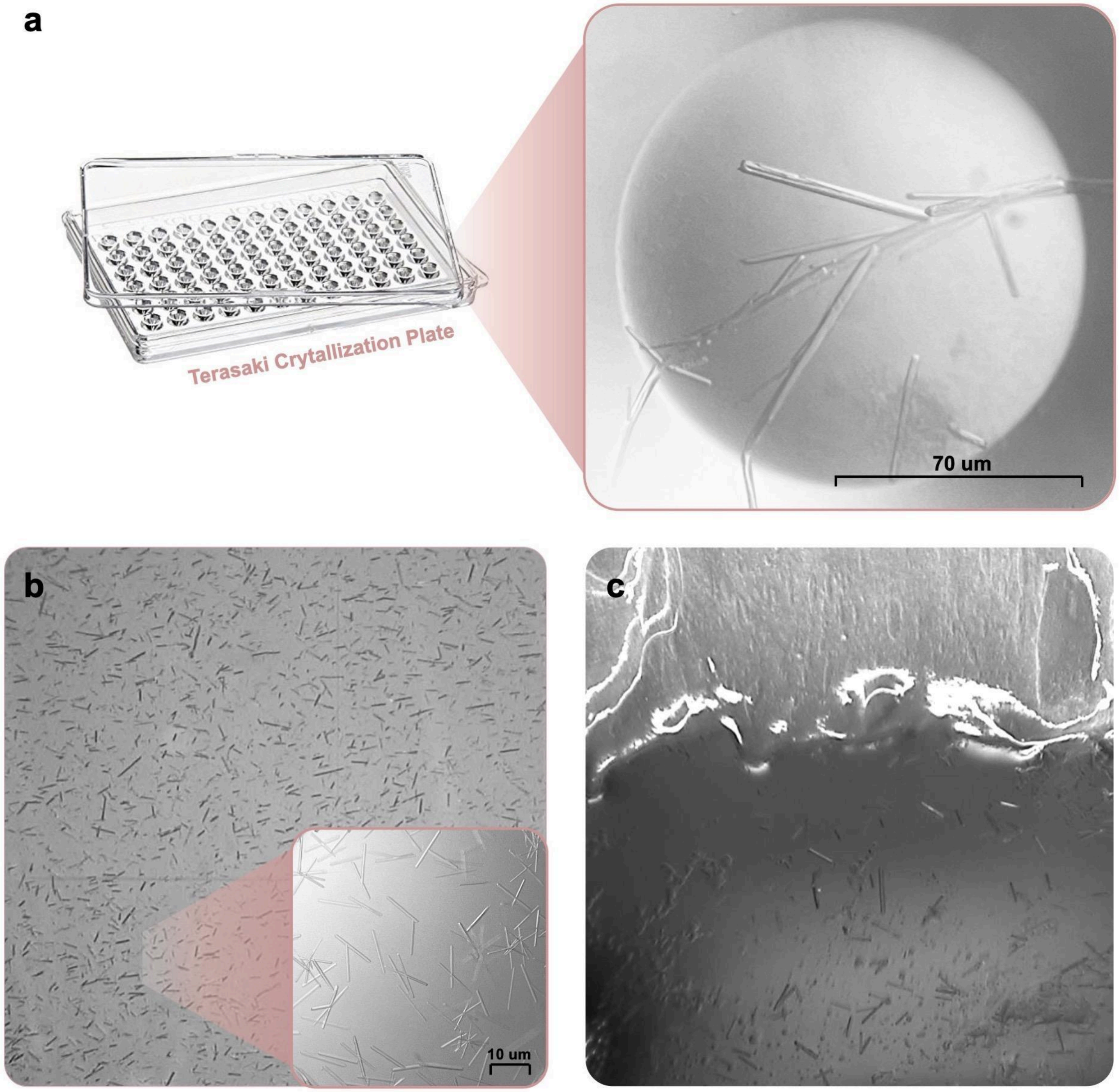


**Supplementary Fig. 2: Crystal images. a** Crystals are obtained from Index™ crystallization screen 1 condition #43 from Hampton Research, USA by using the sitting drop vapor diffusion microbatch technique at room temperature for XRD and synchrotron experiments. **b** Microcrystals are obtained with crystal density $10^8$ by mixing protein and the crystallization condition with a 1:1 ratio at room temperature. **c** Crystal slurry is mixed with grease for sample delivery during SFX experiments.

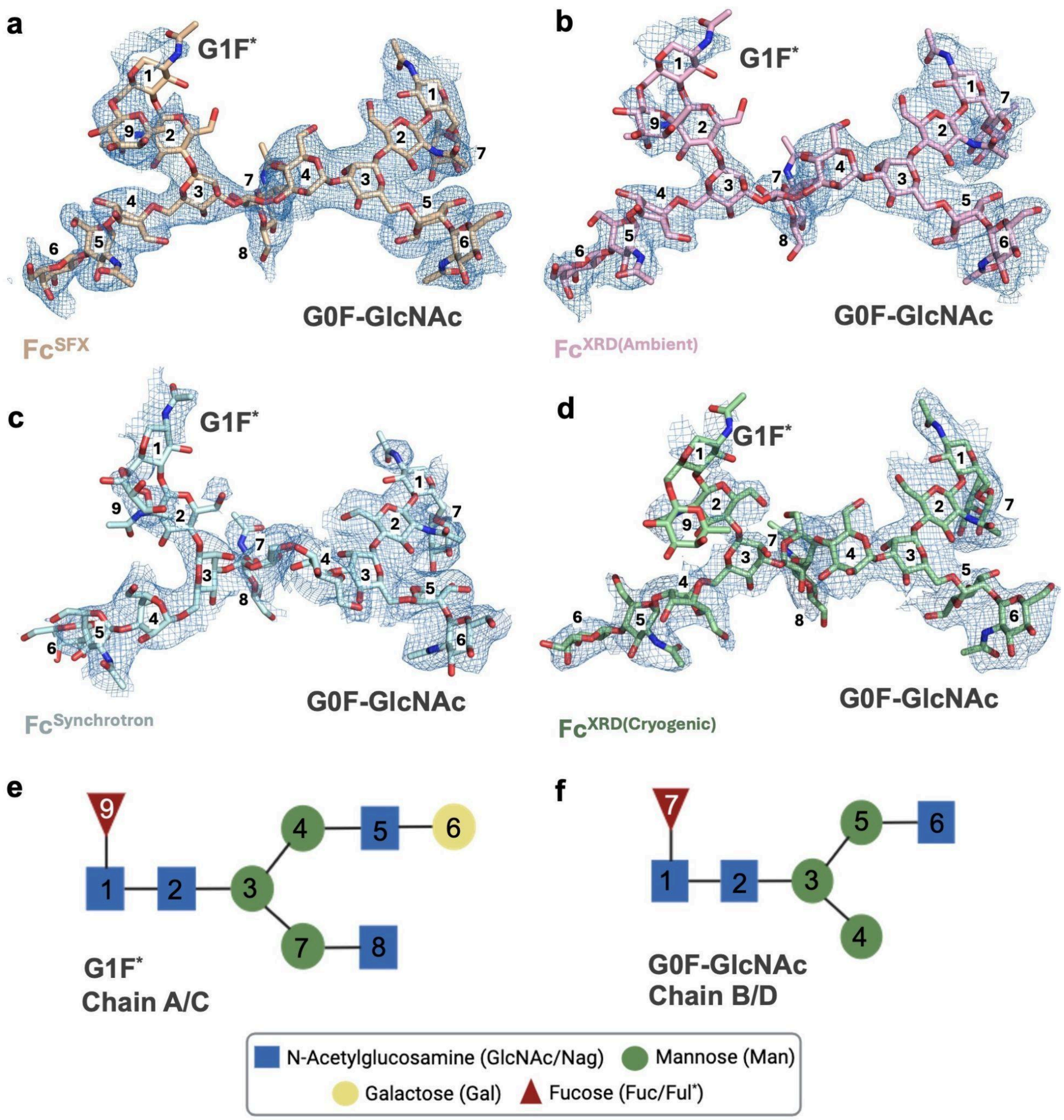


**Supplementary Fig. 3: N-glycan structures for Fc Fragment of IgG1 subtype from biosimilar VEGF-Trap.** The 2Fo-Fc electron densities are contoured at 1 σ level, and colored in skyblue. $Fc^{SFX}$, $Fc^{XRD(Ambient)}$, $Fc^{Synchrotron}$ and $Fc^{XRD(Cryogenic)}$ structures are colored wheat, lightpink, palecyan and palegreen, respectively. PDB IDs and GlyTouCan IDs are indicated in **Supplementary Table 2**.

**a**

| Peptide Fragment | Amino Acid Sequence | Attached PTM | Retetion Time (min) | % Peptide |
|---|---|---|---|---|
| **T32** | EEQYNSTYR | Glycosylation Man5 | 14.15 | 0.67 |
| **T32** | EEQYNSTYR | Glycosylation G1 | 14.91 | 0.65 |
| **T32** | EEQYNSTYR | Glycosylation G0F-GlcNAc | 14.94 | 1.84 |
| **T32** | EEQYNSTYR | Glycosylation G1F* | 15.37 | 38.21 |
| **T32** | EEQYNSTYR | Glycosylation G2F | 15.38 | 4.93 |
| **T32** | EEQYNSTYR | Glycosylation G0F | 15.44 | 50.59 |

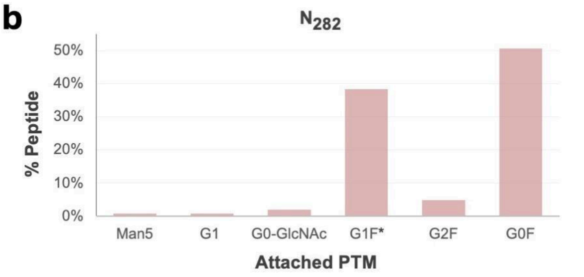


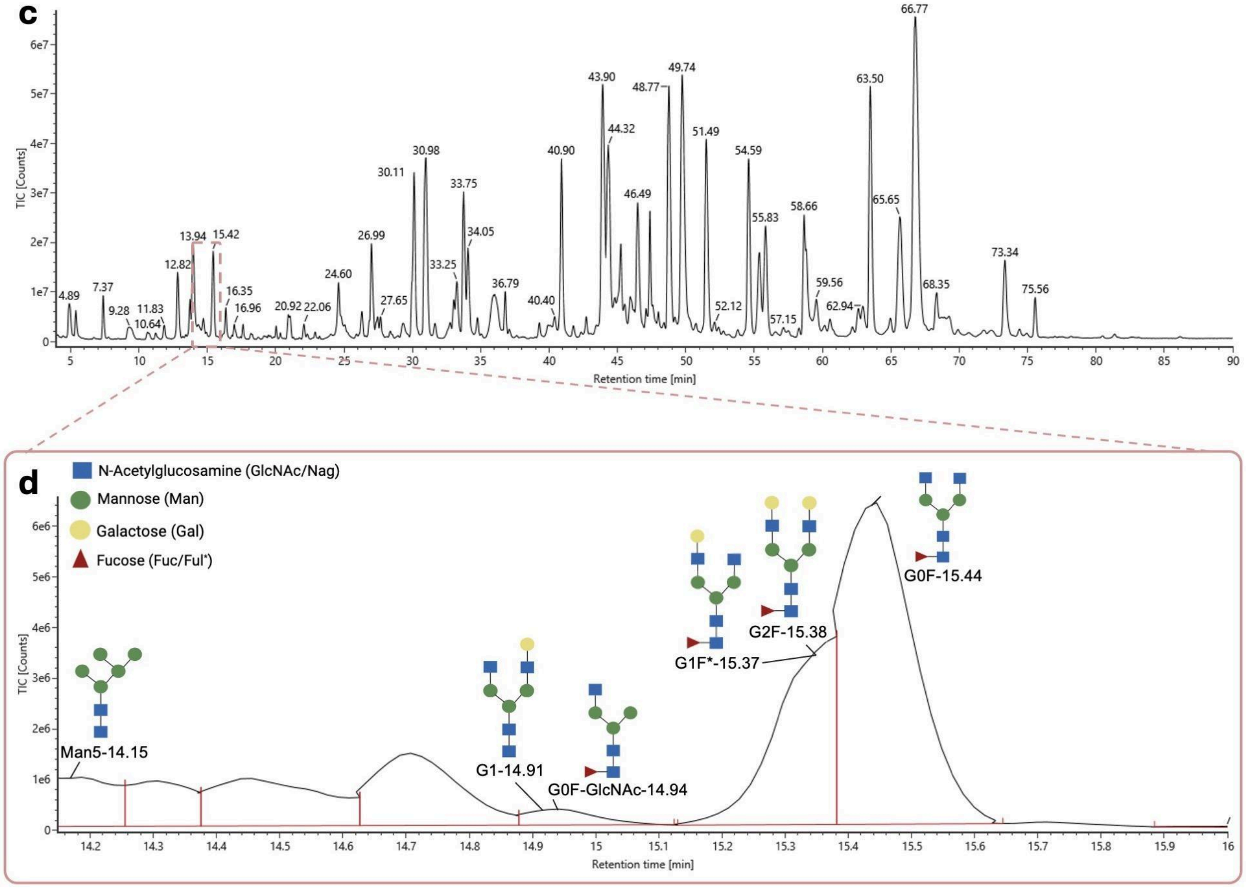


**Supplementary Fig. 4: Mass spectroscopy analysis of the bound glycans to biosimilar VEGF-Trap. a,b** Table and graphs indicate the percentage of glycans that bind to the T32 site ($N_{282}$) in the Fc fragment. **c** Chromatography result for the VEGF-Trap is indicated based on the retention time. **d** The glycans that bound the Fc fragment (T32) are indicated based on the retention time.

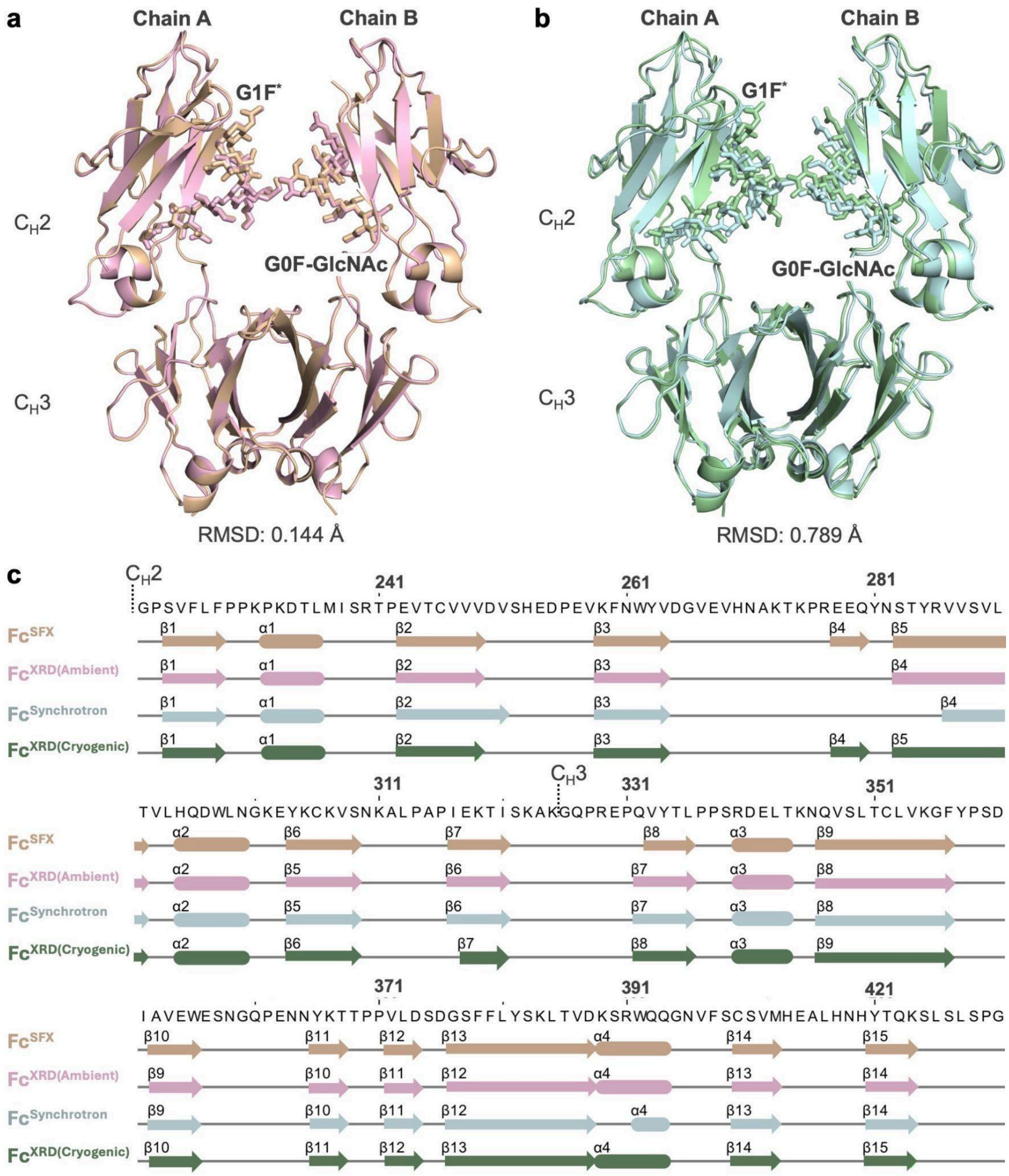


**Supplementary Fig. 5: Structural alignment of the crystal structures of Fc fragment of IgG1 subtype from biosimilar VEGF-Trap. a,b** Ambient ($Fc^{SFX}$ and $Fc^{XRD(Ambient)}$) and cryogenic ($Fc^{Synchrotron}$ and $Fc^{XRD(Cryogenic)}$) temperature structures are aligned and 2D secondary structure alignment is represented in panel **c**. $Fc^{SFX}$, $Fc^{XRD(Ambient)}$, $Fc^{Synchrotron}$ and $Fc^{XRD(Cryogenic)}$ structures are colored wheat, lightpink, palecyan and palegreen, respectively. PDB IDs and GlyTouCan IDs are indicated in **Supplementary Table 2**.

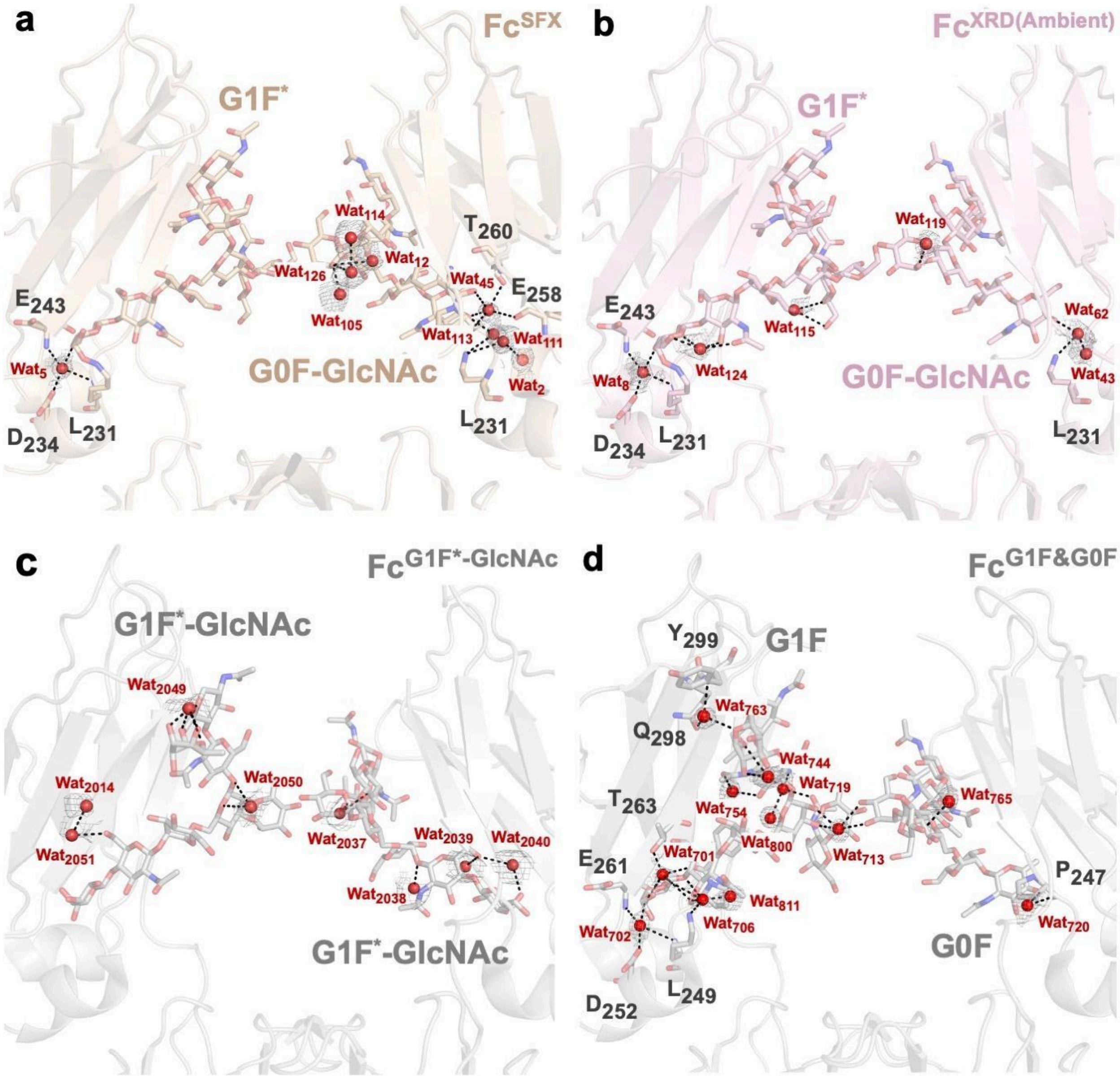


**Supplementary Fig. 6: Water molecules around the glycan-binding pocket.** Water molecules (Wat) are indicated in sphere representation and colored red. Hydrogen bond interactions are shown with dashed black lines. PDB IDs and GlyTouCan IDs are indicated in **Supplementary Table 2**.

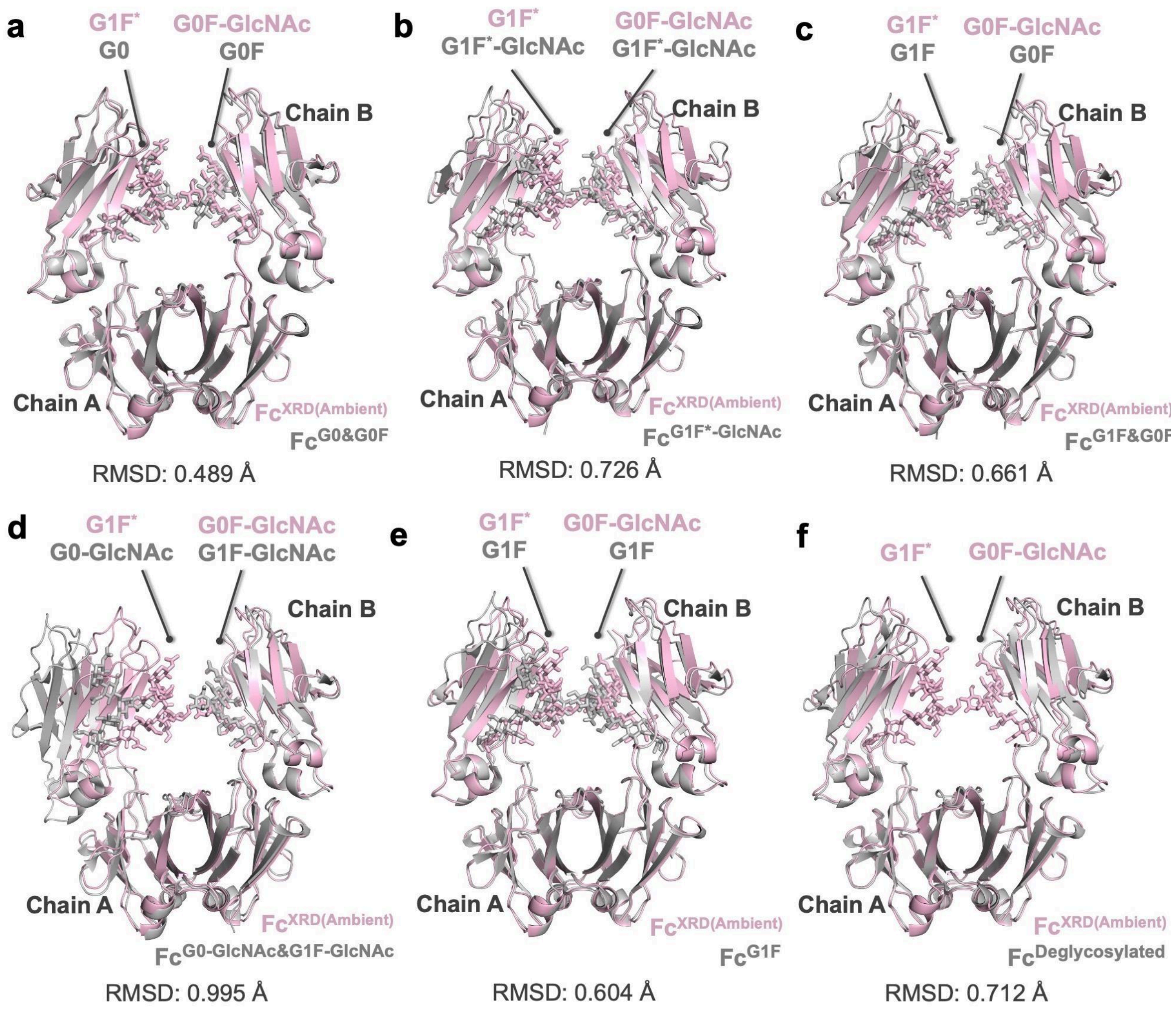


**Supplementary Fig. 7: Conformational changes between $Fc^{XRD(Ambient)}$ and cryogenic temperature structures of the Fc fragment of IgG1 subtype.** $Fc^{XRD}$ structure is superposed with the cryogenic structures of the Fc fragment of IgG1 subtype. $Fc^{XRD}$ structure is colored in wheat while superposed structures are colored in gray. PDB IDs and GlyTouCan IDs are indicated in **Supplementary Table 2**.

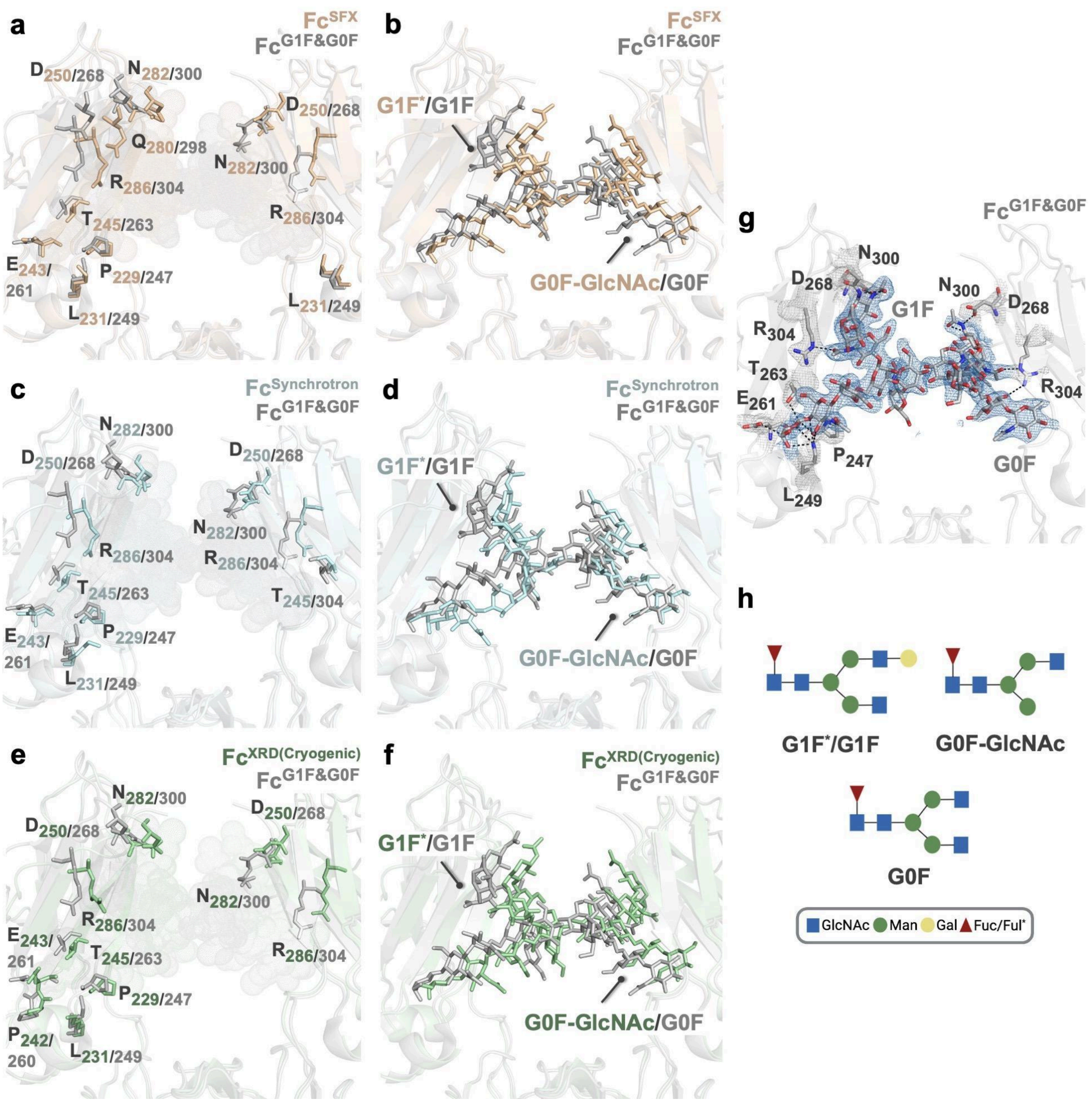


**Supplementary Fig. 8: Comparison of the glycan-binding site of Fc$^{SFX}$, Fc$^{Synchrotron}$ and Fc$^{XRD(Cryogenic)}$ structures with Fc$^{G1F\&G0F}$ structure (PDB ID: 5VGP).** Polar contacts are shown with black dashed lines. The 2Fo-Fc electron densities are contoured at 1 σ level, and colored in skyblue (N-glycans) and gray (protein residues). Fc$^{SFX}$, Fc$^{Synchrotron}$ and Fc$^{XRD(Cryogenic)}$ structures are colored wheat, palecyan and palegreen, respectively while the Fc$^{G1F\&G0F}$ structure is colored gray. PDB IDs and GlyTouCan IDs are indicated in **Supplementary Table 2**.

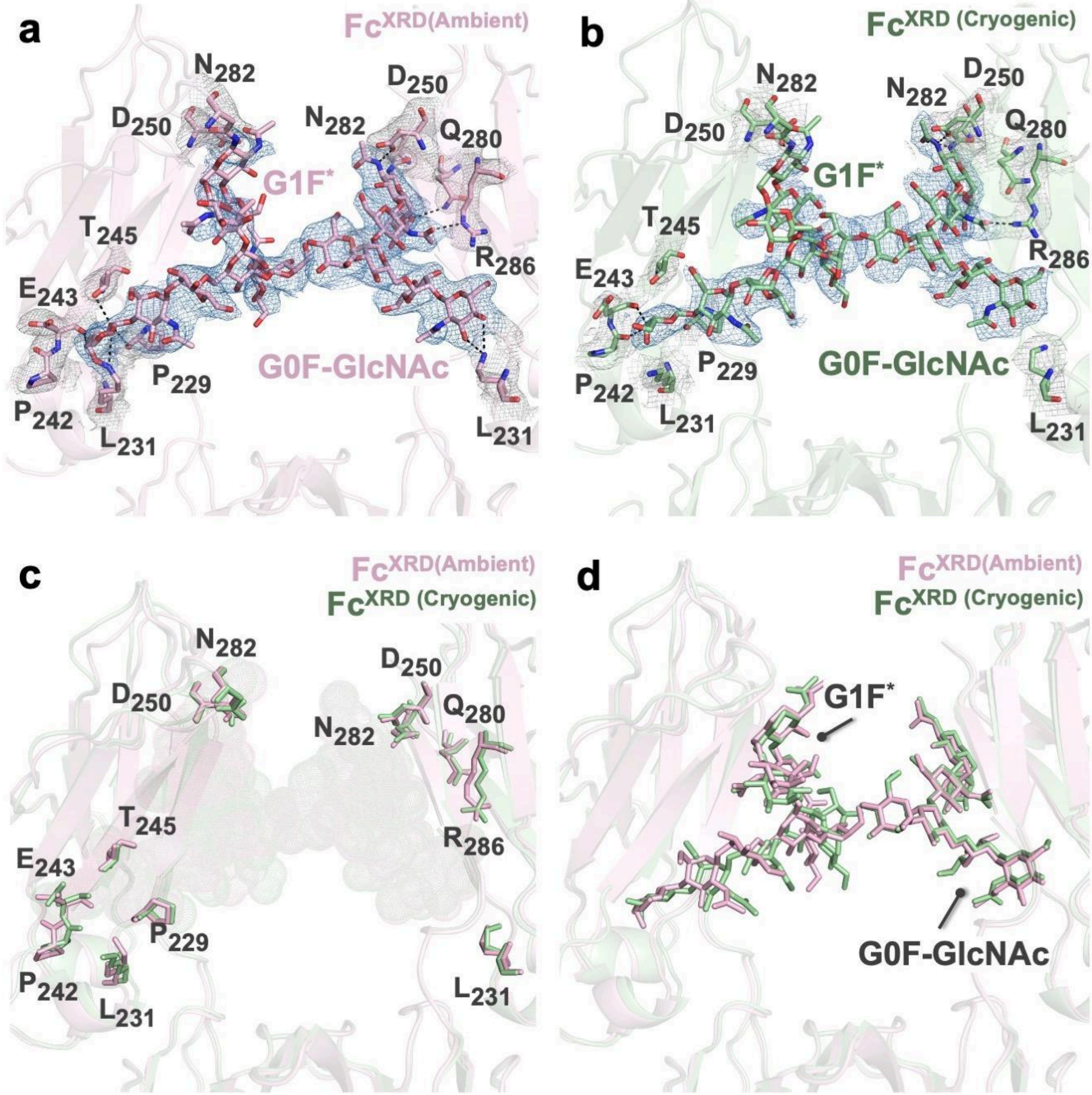


**Supplementary Fig. 9: Conformational changes in the glycan-binding site.** Fc$^{XRD(Ambient)}$ and Fc$^{XRD(Cryogenic)}$ structures are superposed with an RMSD value of 0.451 Å. The 2Fo-Fc electron densities are contoured at 1 σ level, and colored in skyblue (N-glycans) and gray (protein residues). Fc$^{XRD(Ambient)}$and Fc$^{XRD(Cryogenic)}$ are colored lightpink and palegreen, respectively. PDB IDs and GlyTouCan IDs are indicated in **Supplementary Table 2**.

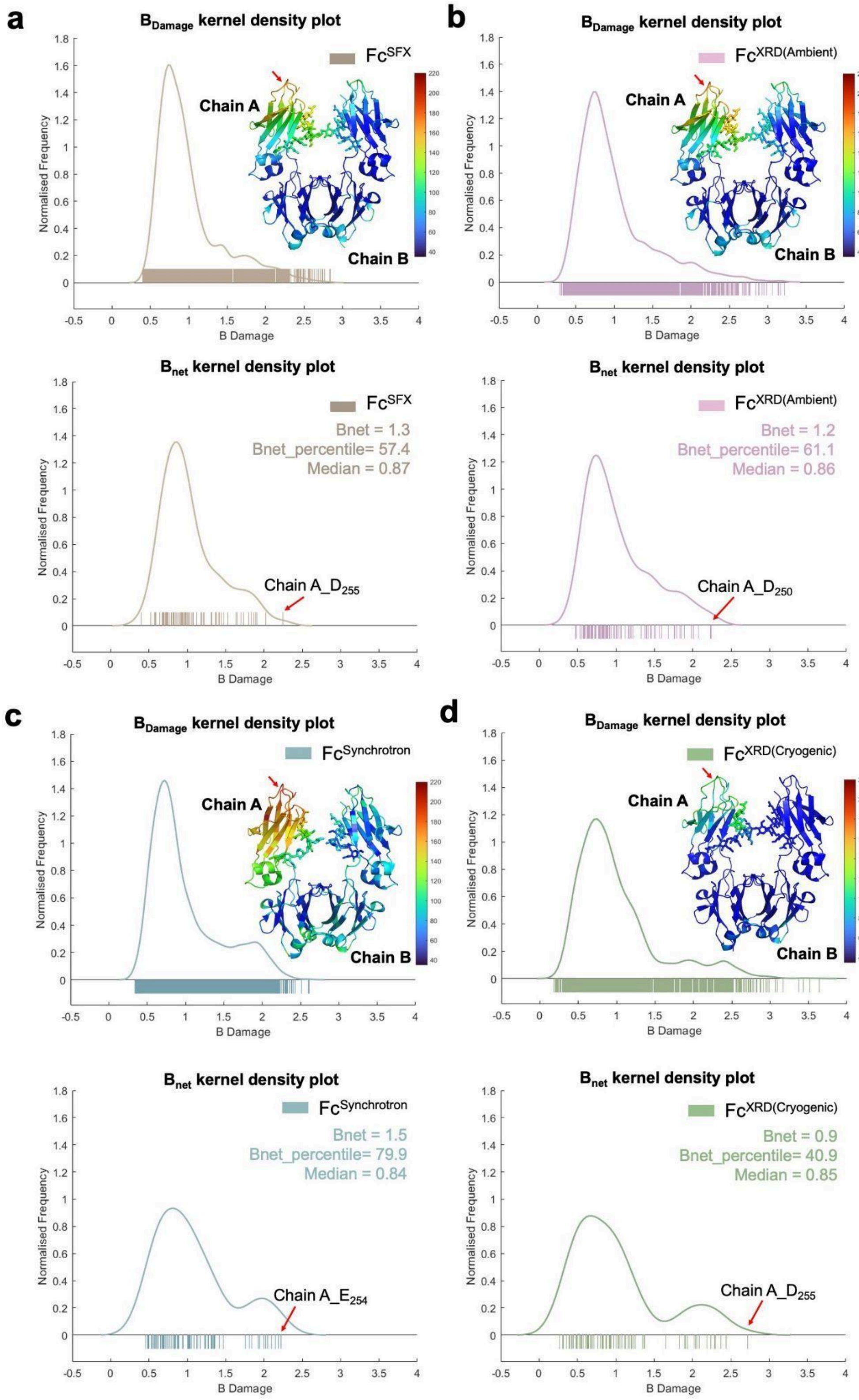


**Supplementary Fig. 10: RABDAM analysis.** Radiation damage for $Fc^{SFX}$, $Fc^{XRD(Ambient)}$, $Fc^{Synchrotron}$ and $Fc^{XRD(Cryogenic)}$ structures are quantified based on atomic B-factor values. $B_{Damage}$ and $B_{net}$ values are represented by kernel density plots. All atoms in the structures are included in the $B_{Damage}$ kernel density plot while only side chain oxygen atoms are considered for the $B_{net}$ kernel density plot. PDB IDs and GlyTouCan IDs are indicated in **Supplementary Table 2**.

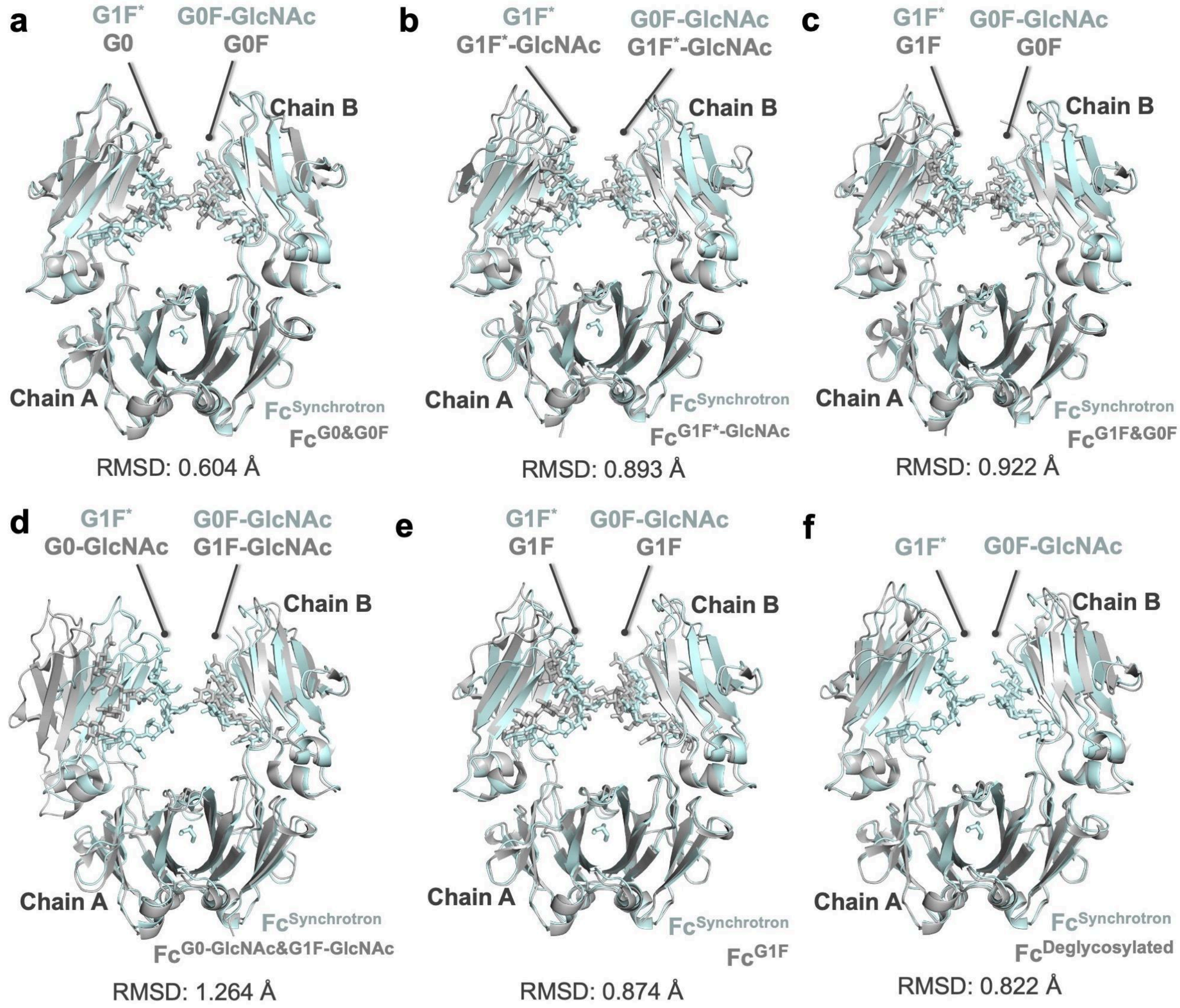


**Supplementary Fig. 11: Conformational changes between $Fc^{Synchrotron}$ and cryogenic temperature structures of Fc fragment of IgG1 subtype.** $Fc^{Synchrotron}$ structure is superposed with the cryogenic structures of the Fc fragment of IgG1. $Fc^{Synchrotron}$ structure is colored in wheat while superposed structures are colored in gray. PDB IDs and GlyTouCan IDs are indicated in **Supplementary Table 2**.

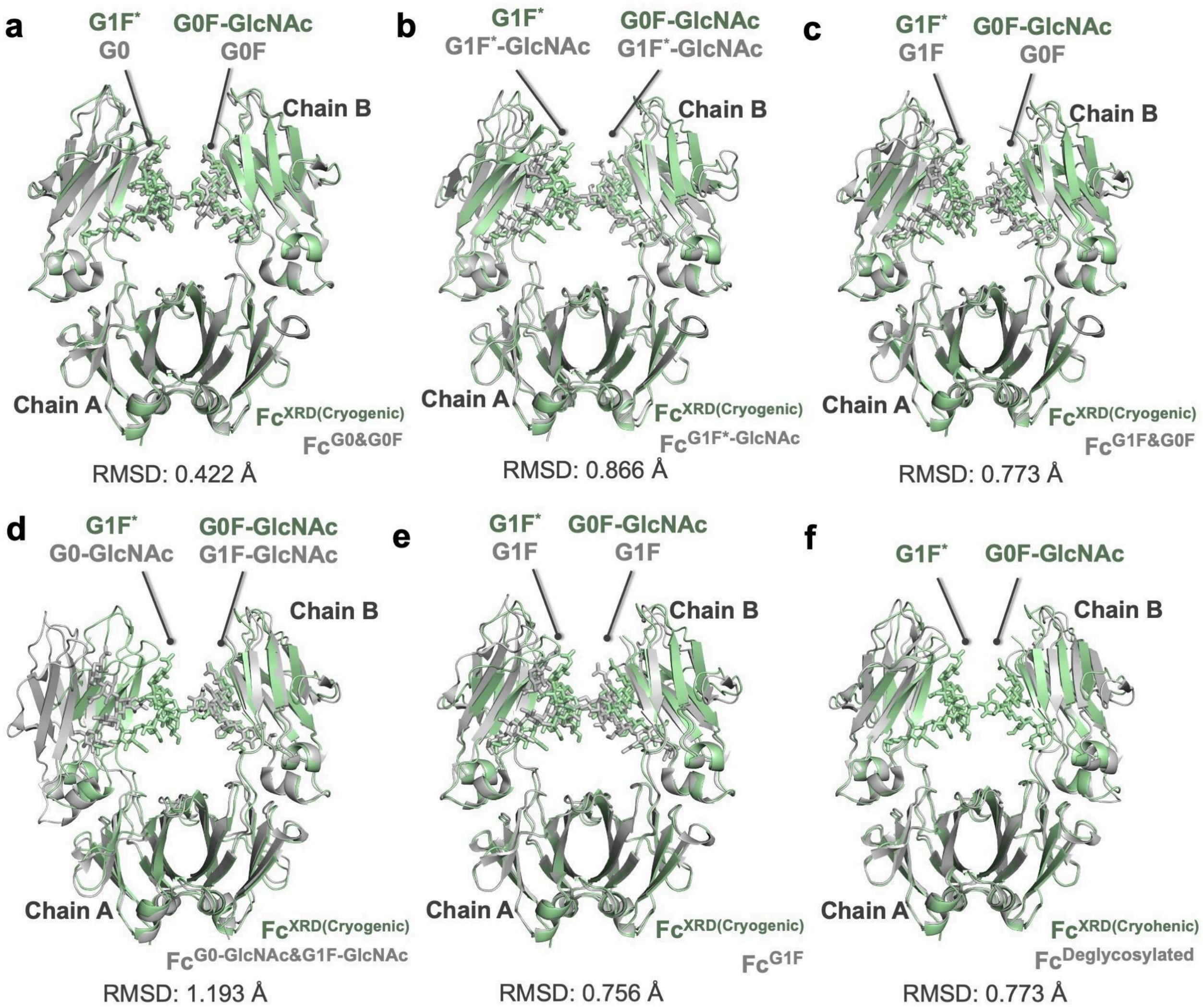


**Supplementary Fig. 12: Conformational changes between $Fc^{XRD(Cryogenic)}$ and cryogenic temperature structures of the Fc fragment of IgG1 subtype.** $Fc^{Synchrotron}$ structure is superposed with the cryogenic structures of the Fc fragment of IgG1. $Fc^{Synchrotron}$ structure is colored in wheat while superposed structures are colored in gray. PDB IDs and GlyTouCan IDs are indicated in **Supplementary Table 2**.

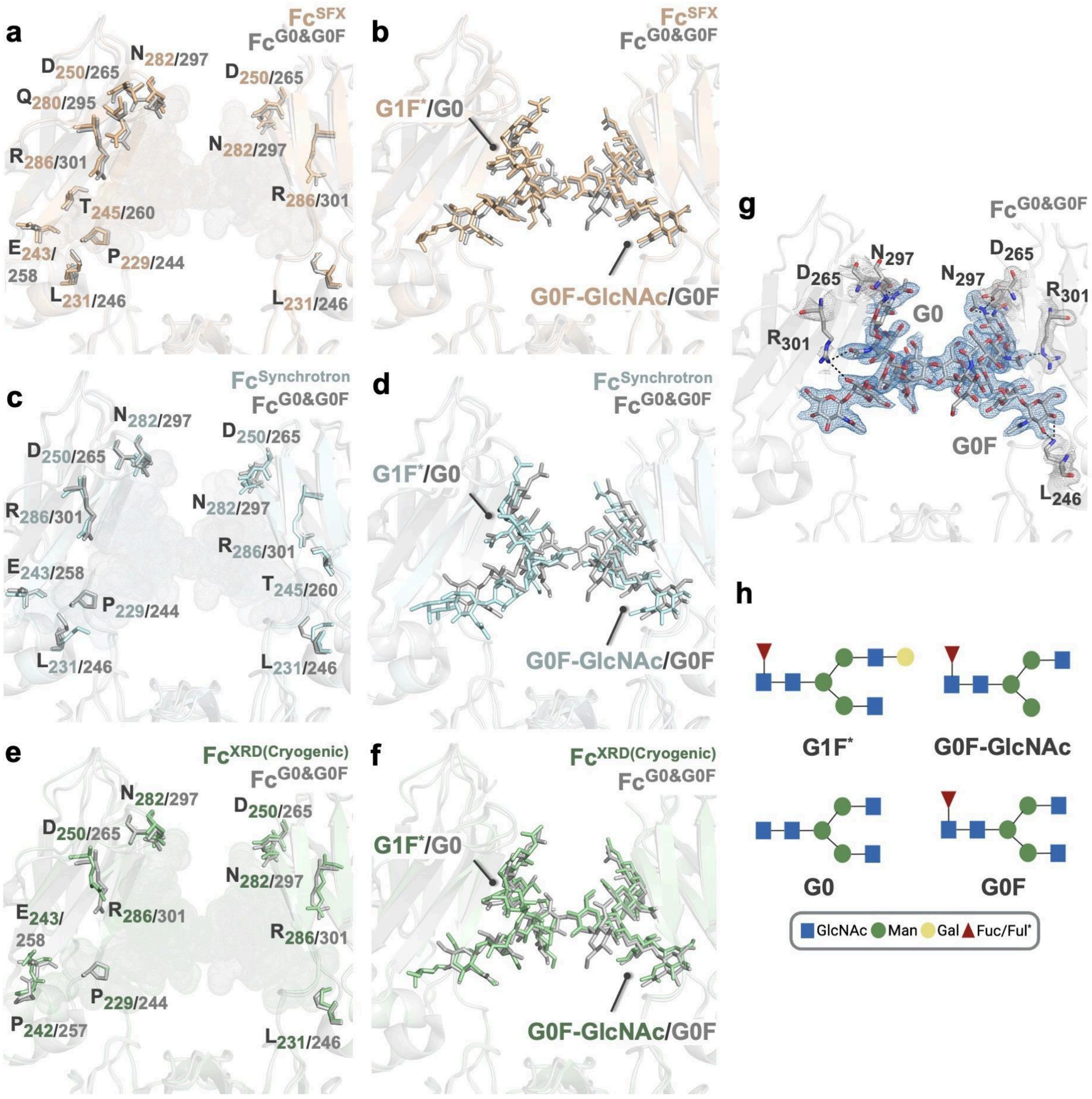


**Supplementary Fig. 13: Comparison of the glycan-binding site of $\mathrm{Fc}^{\mathrm{SFX}}$, $\mathrm{Fc}^{\mathrm{XRD(Cryogenic)}}$ and $\mathrm{Fc}^{\mathrm{Synchrotron}}$ structures with $\mathrm{Fc}^{\mathrm{G0\&G0F}}$ (PDB ID: 4W4N).** Polar contacts are shown with black dashed lines. The 2Fo-Fc electron densities are contoured at 1 σ level, and colored in skyblue (N-glycans) and gray (protein residues). $\mathrm{Fc}^{\mathrm{SFX}}$, $\mathrm{Fc}^{\mathrm{XRD(Cryogenic)}}$ and $\mathrm{Fc}^{\mathrm{Synchrotron}}$ structures are colored wheat, pale green and pale cyan, respectively while the $\mathrm{Fc}^{\mathrm{G0\&G0F}}$ structure is colored gray. PDB IDs and GlyTouCan IDs are indicated in **Supplementary Table 2**.

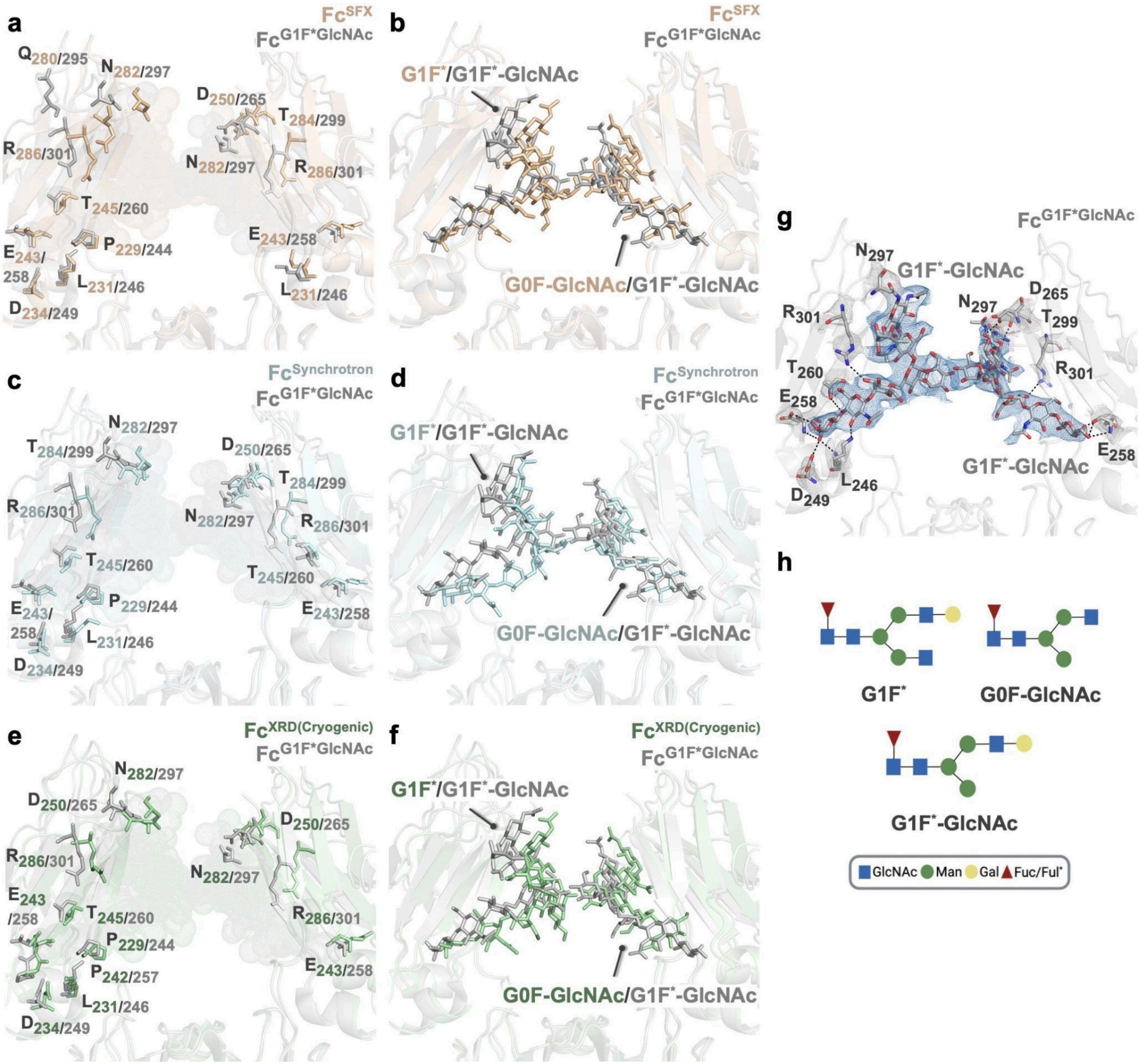


**Supplementary Fig. 14: Comparison of the glycan-binding site of $Fc^{SFX}$, $Fc^{XRD(Cryogenic)}$ and $Fc^{Synchrotron}$ structures with the Fc fragment of IgG1 (PDB ID: 1H3V).** Polar contacts are shown with black dashed lines. The 2Fo-Fc electron densities are contoured at 1 σ level, and colored in skyblue (N-glycans) and gray (protein residues). $Fc^{SFX}$, $Fc^{XRD(Cryogenic)}$ and $Fc^{Synchrotron}$ structures are colored wheat, pale green and pale cyan, respectively while the 1H3V structure is colored gray. GlyTouCan IDs are indicated in **Supplementary Table 2**.

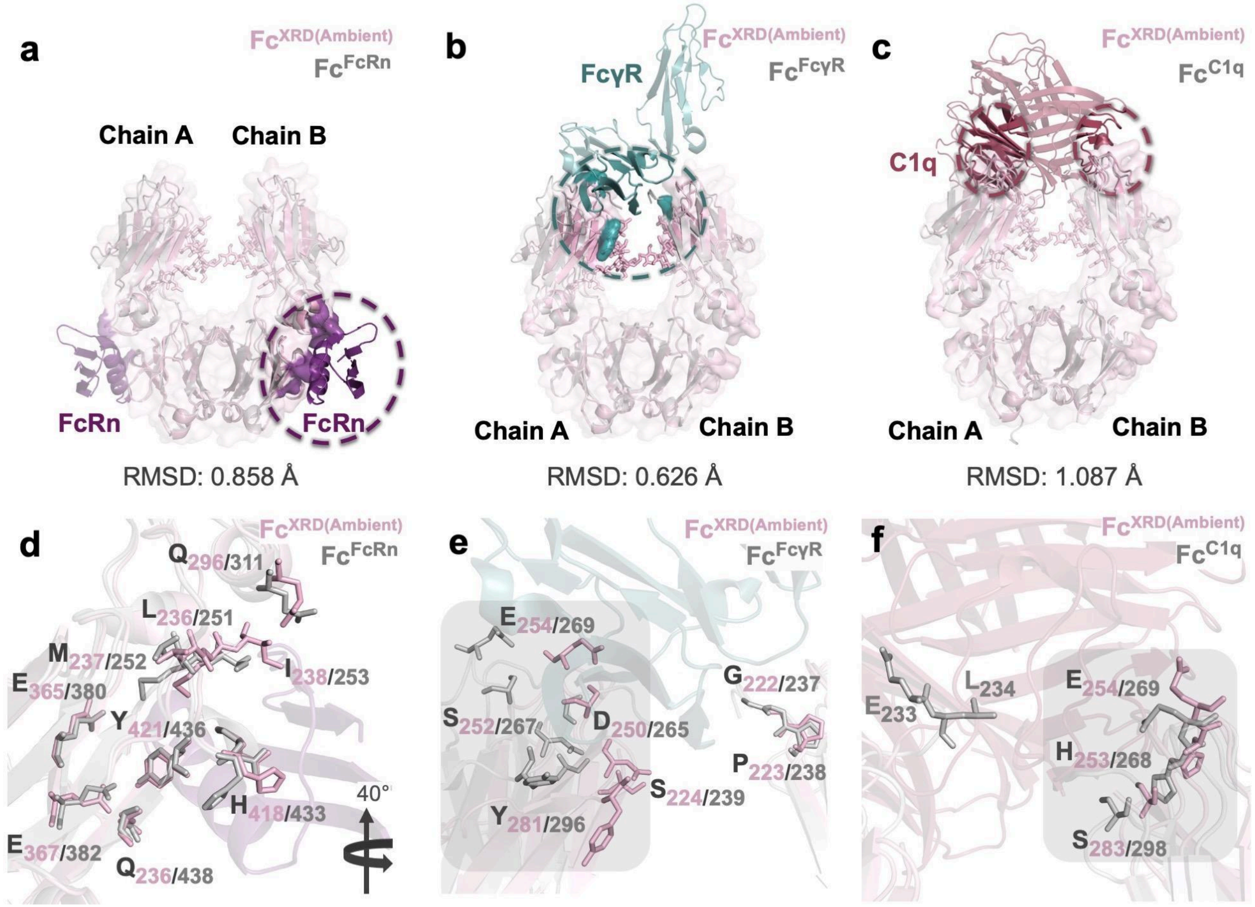


**Supplementary Fig. 15: Conformational changes of the active site residues on Fc fragment in the presence of FcRn, FcγR, and C1q.** Fc$^{XRD(Ambient)}$ structure is colored in lightpink while FcRn, FcγR, and C1q proteins are colored in violetpurple, lightteal and raspberry, respectively. **a,d** Fc fragment of IgG1 subtype in complex with FcRn (Protein G bound) (PDB ID: 1FCC)(Fc$^{FcRn}$) and Fc$^{XRD}$ structures are superposed with RMSD value 0.858 Å. **b,e** Fc fragment of IgG1 subtype in complex with FcγR (PDB ID: 7URU)(Fc$^{Fc\gamma R}$) and Fc$^{XRD}$ structures are superposed with RMSD value 0.626 Å. **c,f** Fc fragment of IgG1 subtype in complex with C1q (PDB ID: 6FCZ)(Fc$^{C1q}$) and Fc$^{XRD}$ structures are superposed with RMSD value 1.087 Å. There is one available binding site for FcγR and 2 potential binding sites for C1q and FcRn per IgG1 molecule. The C1q binding site in Chain A is not visible for Fc$^{XRD}$ and structure as E$_{233}$ and L$_{234}$ residues are located in the hinge region. As a result of superposition, the major shifts in the secondary structures are indicated with gray squares.

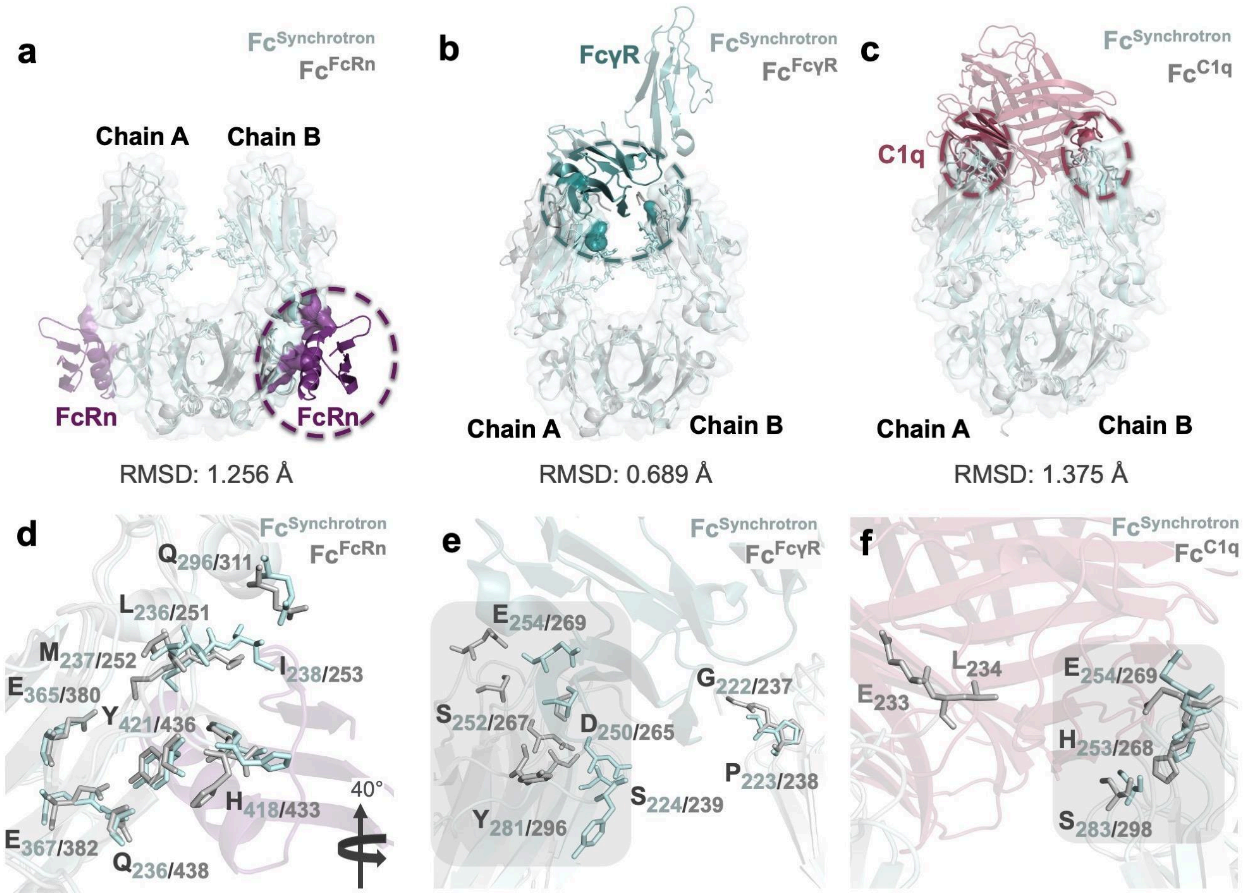


**Supplementary Fig. 16: Conformational changes of the active site residues on the Fc fragment in the presence of FcRn, FcγR and C1q.** $Fc^{Synchrotron}$ structure is colored in lightpink while FcRn, FcγR and C1q proteins are colored in violetpurple, lightteal and raspberry, respectively. **a,d** Fc fragment of IgG1 subtype in complex with FcRn (Protein G bound) (PDB ID: 1FCC)($Fc^{FcRn}$) and $Fc^{Synchrotron}$ structures are superposed with RMSD value 1.256 Å. **b,e** Fc fragment of IgG1 subtype in complex with FcγR (PDB ID: 7URU)($Fc^{FcγR}$) and $Fc^{Synchrotron}$ structures are superposed with RMSD value 0.689 Å. **c,f** Fc fragment of IgG1 subtype in complex with C1q (PDB ID: 6FCZ)($Fc^{C1q}$) and $Fc^{Synchrotron}$ structures are superposed with RMSD value 1.375 Å. There is one available binding site for FcγR and 2 potential binding sites for C1q and FcRn per IgG1 molecule. The C1q binding site in Chain A is not visible for the $Fc^{Synchrotron}$ structure as $E_{233}$ and $L_{234}$ residues are located in the hinge region. As a result of superposition, the major shifts in the secondary structures are indicated with gray squares.

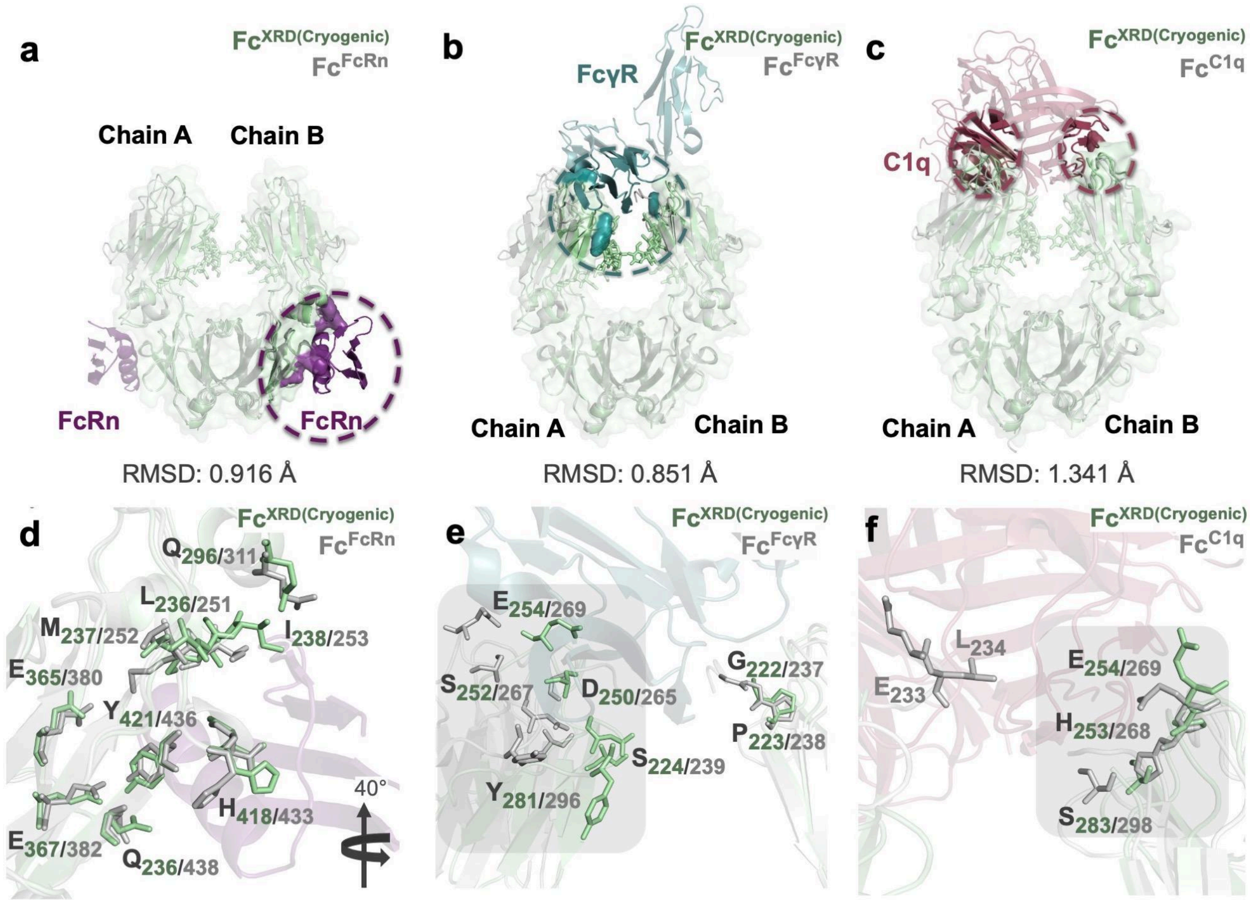


**Supplementary Fig. 17: Conformational changes of the active site residues on Fc fragment in the presence of FcRn, FcγR, and C1q.** $Fc^{XRD(Cryogenic)}$ structure is colored in pale green while FcRn, FcγR, and C1q proteins are colored in violetpurple, lightteal and raspberry, respectively. **a,d** Fc fragment of IgG1 subtype in complex with FcRn (Protein G bound) (PDB ID: 1FCC)($Fc^{FcRn}$) and $Fc^{XRD}$ structures are superposed with RMSD value 0.916 Å. **b,e** Fc fragment of IgG1 subtype in complex with FcγR (PDB ID: 7URU)($Fc^{Fc\gamma R}$) and $Fc^{XRD}$ structures are superposed with RMSD value 0.851 Å. **c,f** Fc fragment of IgG1 subtype in complex with C1q (PDB ID: 6FCZ)($Fc^{C1q}$) and $Fc^{XRD}$ structures are superposed with RMSD value 1.341 Å. There is one available binding site for FcγR and 2 potential binding sites for C1q and FcRn per IgG1 molecule. The C1q binding site in Chain A is not visible for $Fc^{XRD}$ and structure as $E_{233}$ and $L_{234}$ residues are located in the hinge region. As a result of superposition, the major shifts in the secondary structures are indicated with gray squares.

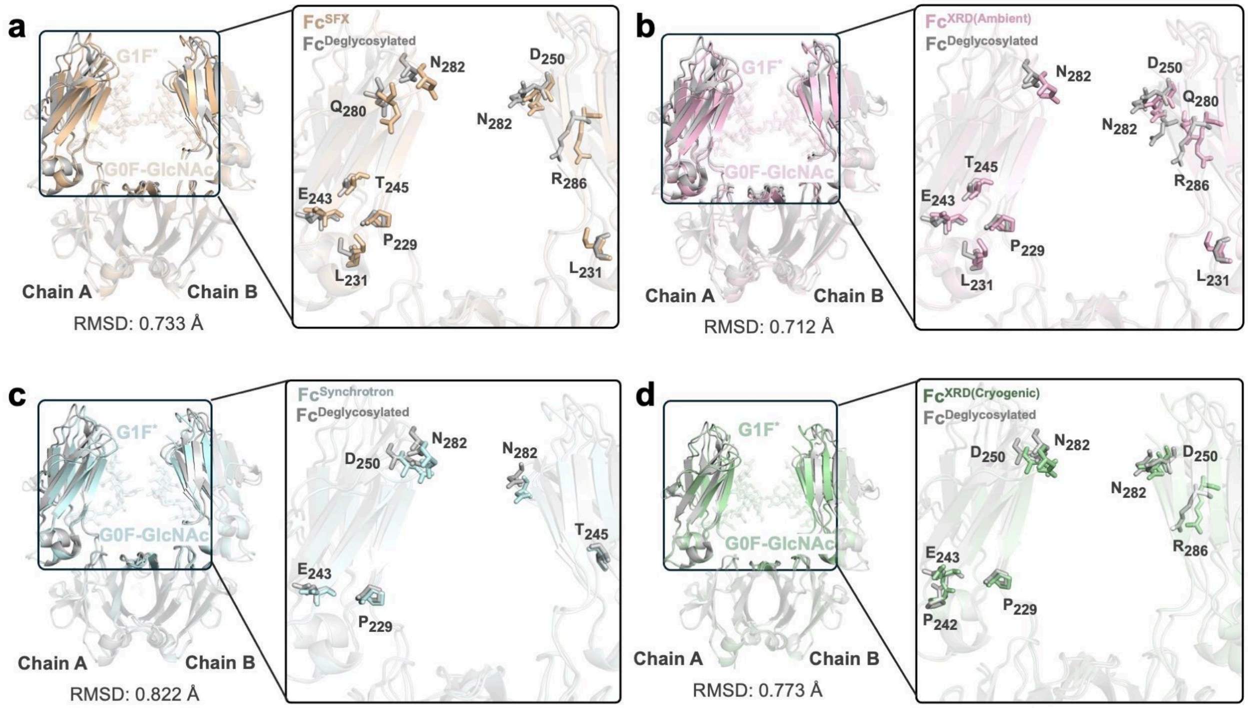


**Supplementary Fig. 18: Conformational changes on the residues that interact with N-glycans (G1F* & G0F-GlcNAc). a** $Fc^{SFX}$ structure is superposed with deglycosylated structure (PDB ID: 7HRO) ($Fc^{Deglycosalated}$) and glycan-binding site residues are indicated in stick representation. The $Fc^{SFX}$ structure is colored in wheat while the $Fc^{Deglycosalated}$ structure is colored gray. **b** $Fc^{XRD(Ambient)}$ structure is superposed with $Fc^{Deglycosalated}$ structure and glycan-binding site residues are indicated in stick representation. $Fc^{XRD(Ambient)}$ structure is colored in lightpink while the $Fc^{Deglycosalated}$ structure is colored gray. **c** $Fc^{Synchrotron}$ structure is superposed with $Fc^{Deglycosalated}$ structure and glycan-binding site residues are indicated in stick representation. The $Fc^{Synchrotron}$ structure is colored in palecyan while the $Fc^{Deglycosalated}$ structure is colored gray. **d** $Fc^{XRD(Cryogenic)}$ structure is superposed with $Fc^{Deglycosalated}$ structure and glycan-binding site residues are indicated in stick representation. The $Fc^{XRD(Cryogenic)}$ structure is colored pale green while the $Fc^{Deglycosalated}$ structure is colored gray. PDB IDs and GlyTouCan IDs are indicated in **Supplementary Table 2**.

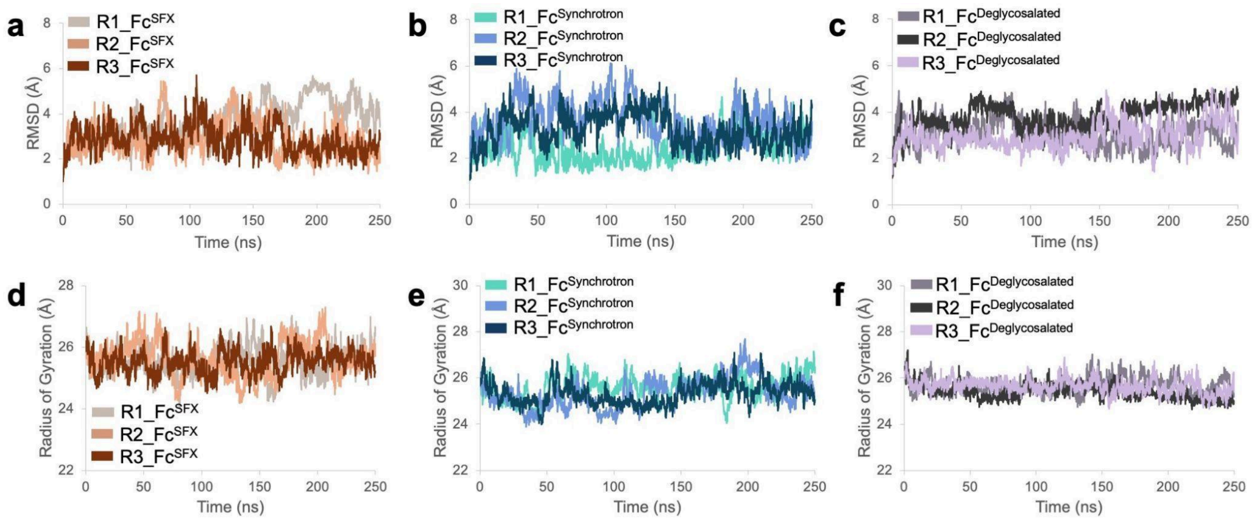


**Supplementary Fig. 19: RMSD (Å) and Radius of Gyration (Å) calculations for each production run.** RMSD (Å) and Radius of Gyration (Å) for each protein structure are calculated based on the backbone through 250 ns simulation. Each production run for Fc$^{SFX}$, Fc$^{Synchrotron,}$ and Fc$^{Deglycosalated}$ is symbolized as R1, R2, and R3.

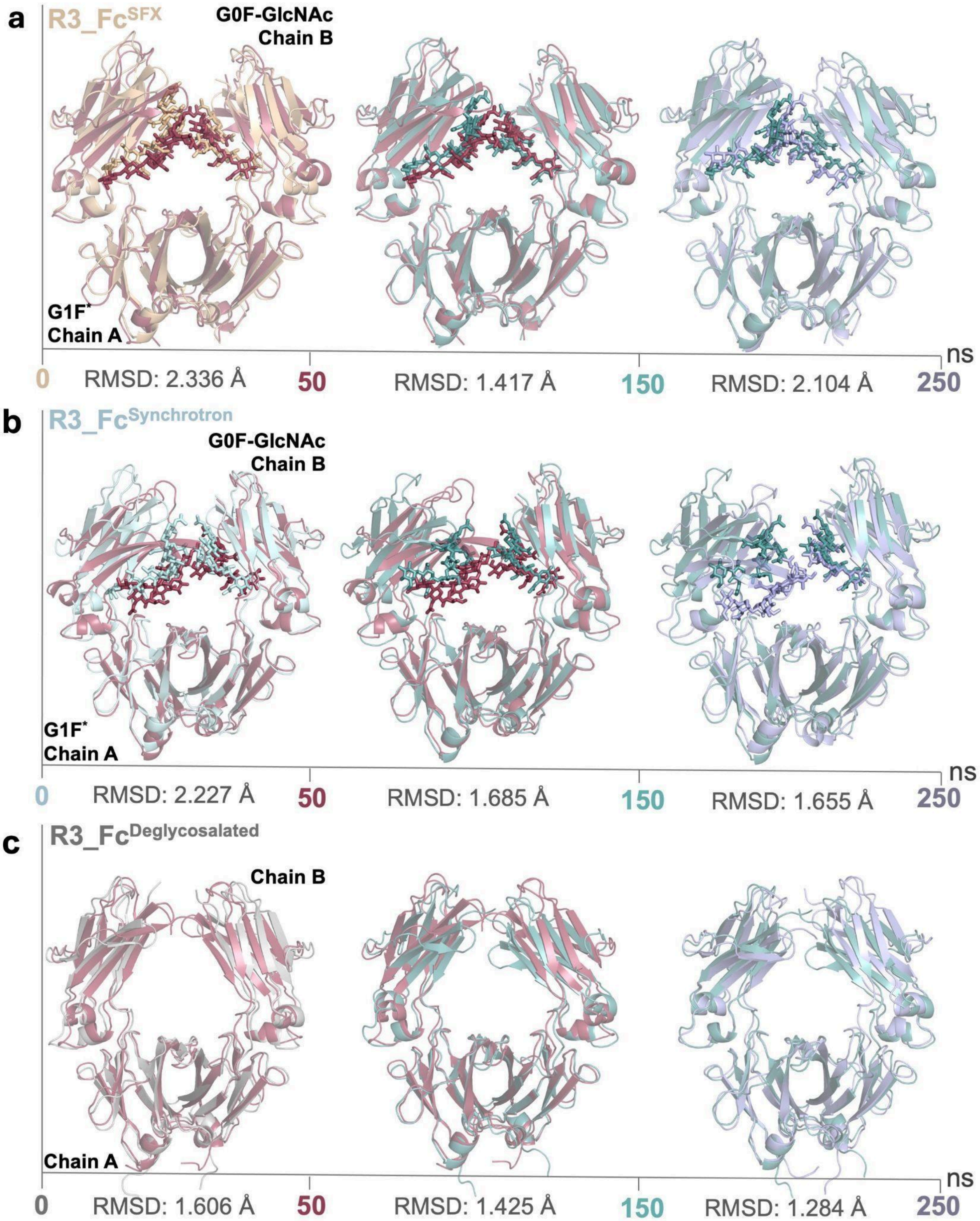


**Supplementary Fig. 20: Conformational changes on Fc fragment through 250 ns Molecular Dynamic simulation.** To analyze conformational changes, the coordinates in the corresponding frame to time are saved as a .pdb file and snapshots are visualized with cartoon (protein) and stick (N-glycans) representation. Three time points are used from the first production run (R3) of simulations for each structure.

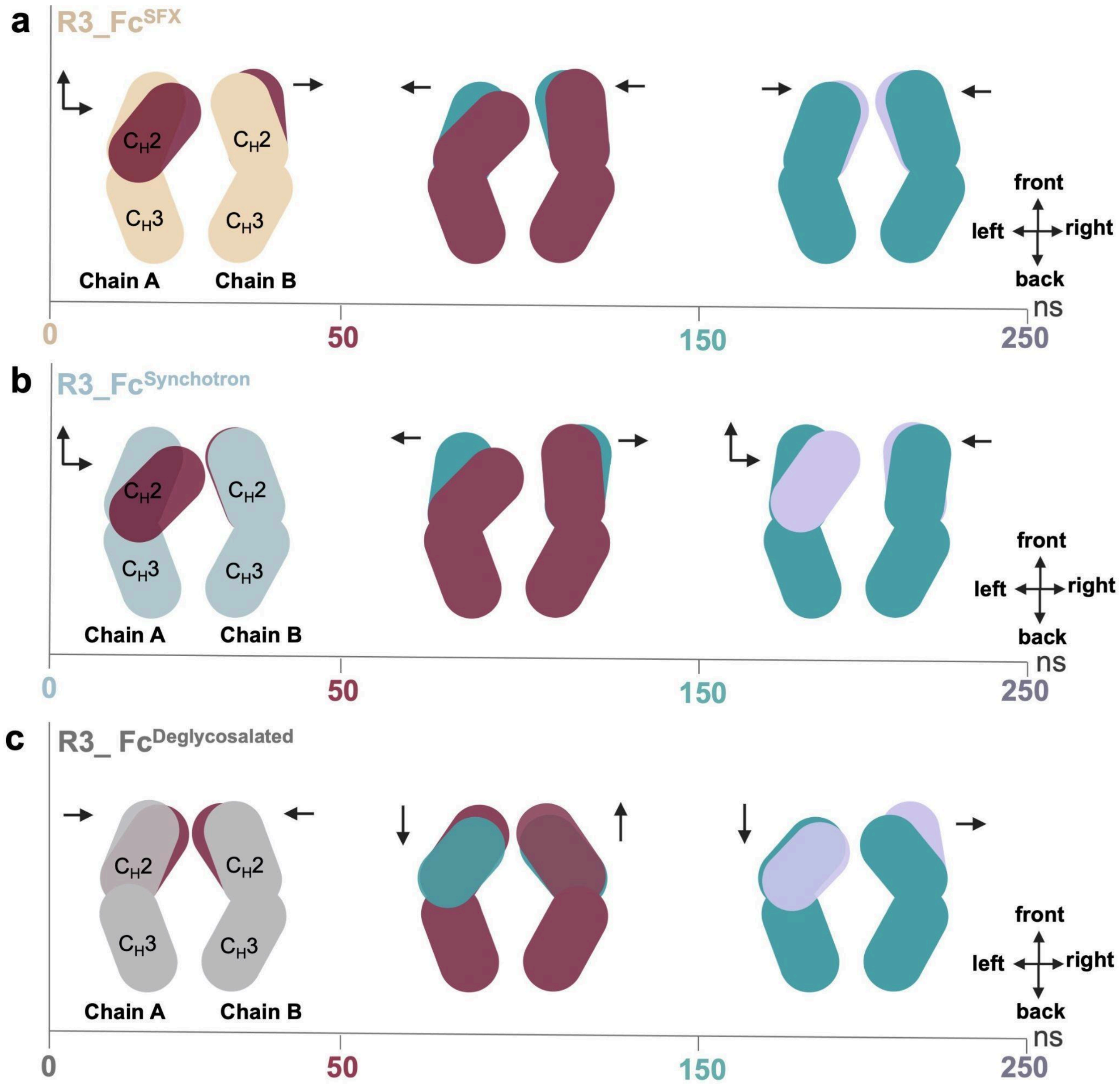


**Supplementary Fig. 21: Systematic representation of the movements of chains through 250 ns Molecular Dynamic simulation.** The figure is generated based on **Supplementary Figure 20.** The direction of the movement for the $C_H2$ domain is indicated with black arrows.

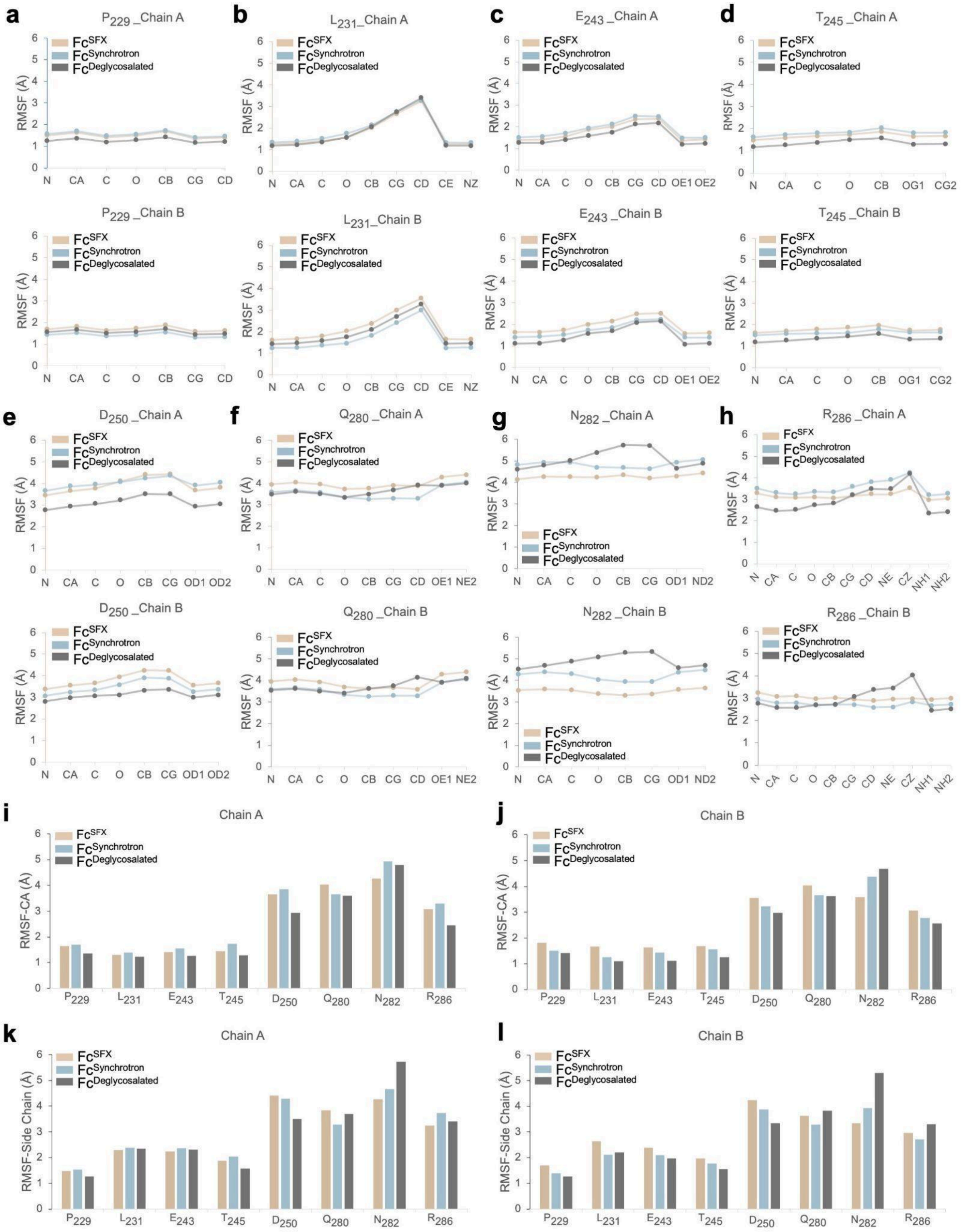
a
P229 _Chain A
P229 _Chain B
b
L231_Chain A
L231_Chain B
c
E243 _Chain A
E243 _Chain B
d
T245 _Chain A
T245 _Chain B
e
D250 _Chain A
D250 _Chain B
f
Q280 _Chain A
Q280 _Chain B
g
N282 _Chain A
N282 _Chain B
h
R286 _Chain A
R286 _Chain B
Fc^SFX
Fc^Synchrotron
Fc^Deglycosalated
RMSF (Å)
N CA C O CB CG CD CE NZ OE1 OE2 OG1 CG2 OD1 OD2 NE2 ND2 NE CZ NH1 NH2
i
Chain A
j
Chain B
RMSF-CA (Å)
k
Chain A
l
Chain B
RMSF-Side Chain (Å)
P229 L231 E243 T245 D250 Q280 N282 R286

**Supplementary Fig. 22: RMSF calculation for the critical residues during N-glycosylation. (a-h)** Atom versus RMSF (Å) is calculated for each residue that performs polar contact with N-glycans based on crystal structures. **I,j** RMSF (Å) based on alpha carbon (CA) atom is calculated for each residue. **k,l** RMSF (Å) based on side chain carbon atoms is calculated for each residue. RMSF (Å) values of the side chain carbon atoms for each residue are averaged. The average calculation was not performed for $T_{245}$ as if it has only one side chain carbon atom. The standard error of the mean (SEM) is indicated in **Supplementary Table 3**.

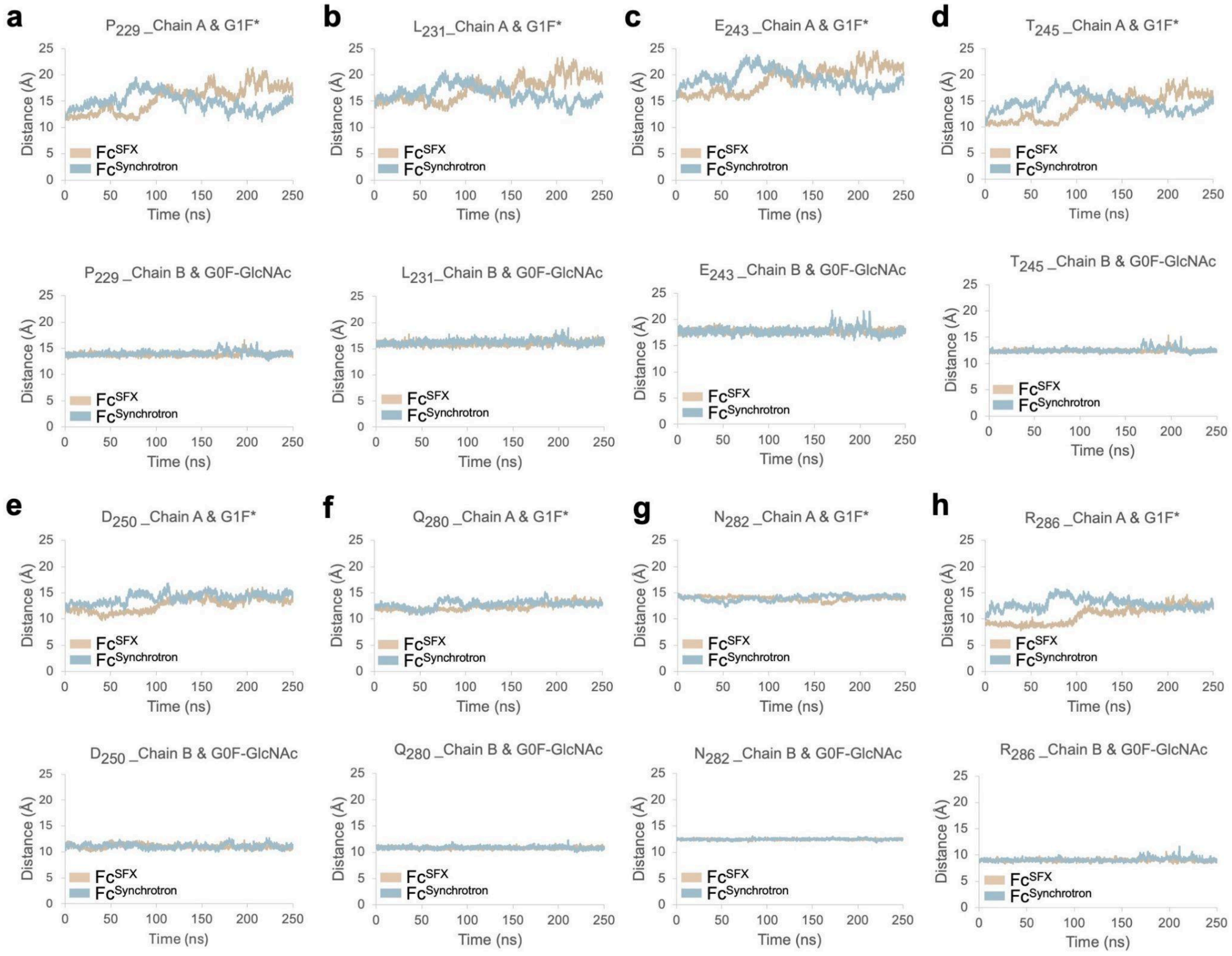


**Supplementary Fig. 23: Distance between critical residues and N-glycans (G1F* and G0F-GlcNAc) during 250 ns Molecular Dynamic simulation.** Residue-glycan distances based on the center of mass were calculated for $Fc^{SFX}$ and $Fc^{Synchrotron}$ structures.

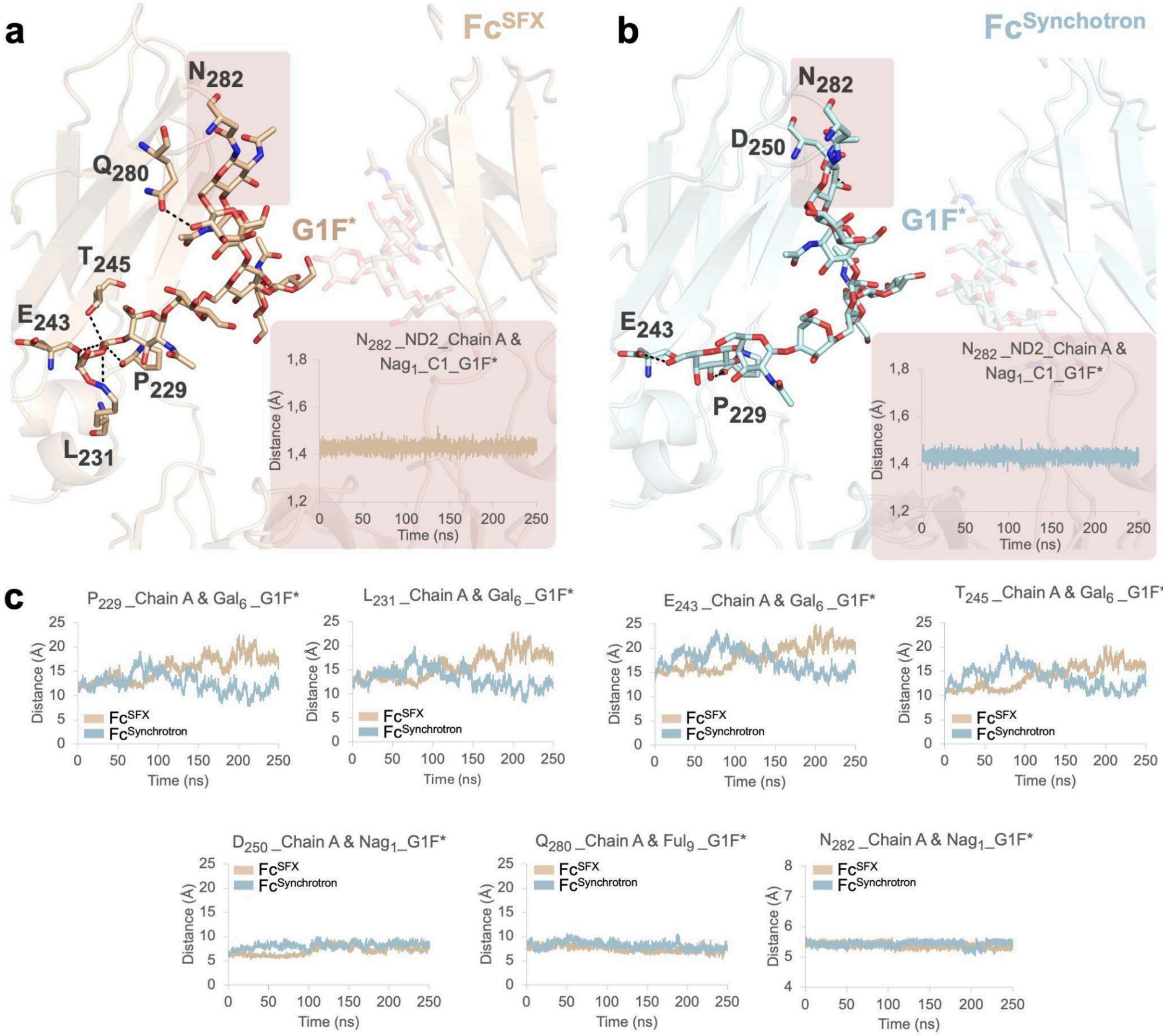


**Supplementary Fig. 24: Distance between critical residues and N-glycan (G1F*) during 250 ns Molecular Dynamic simulation. a,b** The atom-atom distance for $N_{282}$ and G1F* is calculated to indicate the stability of N-glycosylation through simulation. **c** Residue-residue distances based on the center of mass are calculated for critical residues.

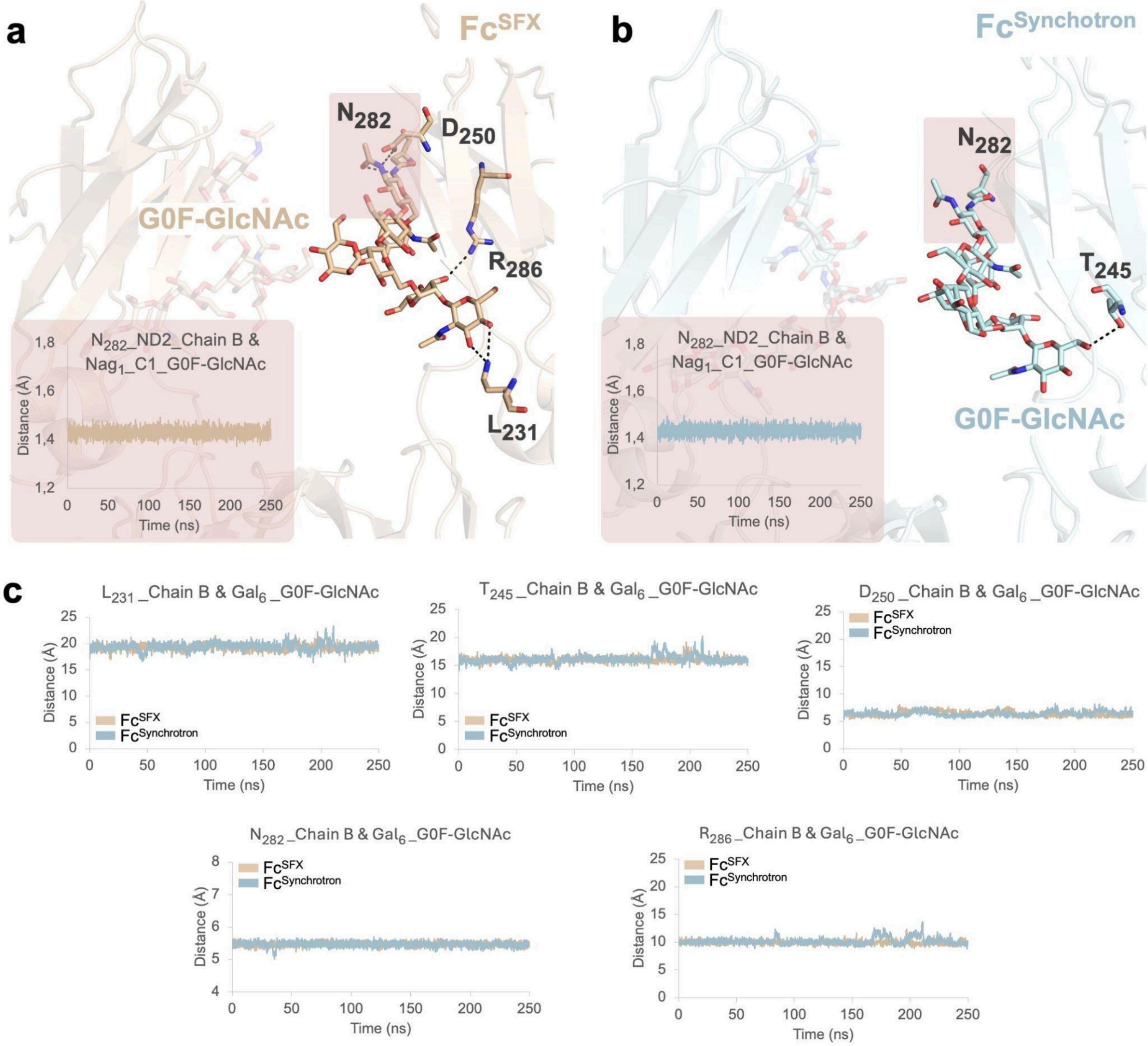


**Supplementary Fig. 25: Distance between critical residues and N-glycan (G0F-GlcNAc) during 250 ns Molecular Dynamic simulation. a,b** The atom-atom distance for $N_{282}$ and G0F-GlcNAc is calculated to indicate the stability of N-glycosylation through simulation. **c** Residue-residue distances based on the center of mass are calculated for critical residues.

**Supplementary Table 1: Data collection and structure refinement statistics.** The highest resolution shell is shown in parentheses.

| Datasets | $Fc^{SFX}$ | $Fc^{XRD(Ambient)}$ | $Fc^{Synchrotron}$ | $Fc^{XRD(Cryogenic)}$ |
|---|---|---|---|---|
| **PDB IDs** | 8ZCK | 8ZCL | 8ZCM | 9IIE |
| **Data Collection temperature** | Ambient | Ambient | Cryogenic | Cryogenic |
| **Data collection** | | | | |
| **X-ray source** | *SACLA (BL2)* | *Turkish DeLight* | *Spring-8 (BL32XU)* | *Turkish DeLight* |
| **Space group** | $P\ 2_12_12_1$ | $P\ 2_12_12_1$ | $P\ 2_12_12_1$ | $P\ 2_12_12_1$ |
| ***a, b, c*** **(Å)** | 50.47, 79.99, 143.19 | 50.47, 79.99, 143.19 | 49.45, 79.63, 125.54 | 50.00, 80.69, 136.79 |
| **α, β, γ (°)** | 90, 90, 90 | 90, 90, 90 | 90, 90, 90 | 89.99,89.94,89.93 |
| **Resolution (Å)** | 32.67-2.00 (2.14-2.00) | 143.31-2.00 (2.05-2.00) | 46.01-2.64 (2.80-2.64) | 137.08-3.14 (3.22-3.14) |
| $I / \sigma I$ | 4.49(2.06) | 5.73(0.01) | 7.55(0.97) | 2.26 (0.75) |
| **[1]*CC** or [2]*CC(1/2)*** | [1]0.99(0.41) | [1]0.96(0.29) | [2]99.3(44.1) | [1]0.92(0.51) |
| **Completeness (%)** | 100.00(100.00) | 96.9(85.00) | 95.5(97.4) | 96.5 (98.6) |
| **Redundancy** | 245.24(170.5) | 38.5(2.8) | 2.85(2.77) | 3.6 (3.9) |
| **Refinement** | | | | |
| **Resolution (Å)** | 29.20-2.00 (2.13-2.00) | 40.99-2.60 (2.70-2.60) | 46.01-2.64 (2.73-2.64) | 23.84-3.14 (3.34-3.14) |
| **No. reflections** | 35449 | 17036 | 14530 | 8053 |
| $R_{work} / R_{free}$ | 0.20/0.23 (0.41/0.37) | 0.17/0.23 (0.34/0.40) | 0.35/0.40 (0.36/0.39) | 0.25/0.30 (0.34/0.41) |
| **No. atoms** | | | | |
| **Protein** | 3348 | 3330 | 3335 | 3318 |
| **Ligand/ion/Water** | 322 | 285 | 224 | 225 |
| ***B*-factors** | | | | |
| **Protein** | 68.97 | 63.89 | 90.40 | 42.54 |
| **Ligand/ion/Water** | 83.20 | 85.78 | 85.87 | 48.23 |
| **R.m.s. deviations** | | | | |
| **Bond lengths (Å)** | 0.003 | 0.004 | 0.003 | 0.003 |
| **Bond angles (°)** | 0.553 | 0.602 | 0.514 | 0.503 |
| **Ramachandran plot** | | | | |
| **Favored (%)** | 99.27 | 99.27 | 98.78 | 96.83 |
| **Allowed (%)** | 0.73 | 0.73 | 1.22 | 3.17 |
| **Disallowed (%)** | 0.00 | 0.00 | 0.00 | 0.00 |

**Supplementary Table 2: GlyTouCan IDs.** GlyTouCan IDs are indicated based on the N-glycan type for each structure.

| | | | | | |
|---|---|---|---|---|---|
| **$\text{Fc}^{\text{SFX}}$ ,$\text{Fc}^{\text{XRD(Ambient)}}$, $\text{Fc}^{\text{Synchrotron}}$ and $\text{Fc}^{\text{XRD(Cryogenic)}}$** | **PDB IDs: 8ZCK, 8ZCL, 8ZCM and 9IIE,respectively** | | **$\text{Fc}^{\text{G0\&G0F}}$** | **PDB ID: 4W4N** | |
| **N-glycan Type** | G1F* | G0F-GlcNAc | **N-glycan Type** | G0 | G0F |
| **GlyTouCan ID** | G56907CZ | G45889JQ | **GlyTouCan ID** | G39213VZ | G80858MF |
| **$\text{Fc}^{\text{G1F*-GlcNAc}}$** | **PDB ID: 1H3V** | | **$\text{Fc}^{\text{G1F\&G0F}}$** | **PDB ID: 5VGP** | |
| **N-glycan Type** | G1F*-GlcNAc | G1F*-GlcNAc | **N-glycan Type** | G1F | G0F |
| **GlyTouCan ID** | G82081LV | G83535HI | **GlyTouCan ID** | G27919IH | G80858MF |
| **$\text{Fc}^{\text{G0-GlcNAc\&G1F-GlcNAc}}$** | **PDB ID: 5JII** | | **$\text{Fc}^{\text{G1F}}$** | **PDB ID: 7LBL** | |
| **N-glycan Type** | G0-GlcNAc | G1F-FlcNAc | **N-glycan Type** | G1F | G1F |
| **GlyTouCan ID** | G64481DJ | G661334IA | **GlyTouCan ID** | G27919IH | G27919IH |

**Supplementary Table 3:** The standard error of the mean (SEM) values for Supplementary Fig. 22k,l.

| Chain A | | | | | | | |
|---|---|---|---|---|---|---|---|
| **SEM** | $P_{229}$ | $L_{231}$ | $E_{243}$ | $D_{250}$ | $Q_{280}$ | $N_{282}$ | $R_{286}$ |
| $Fc^{SFX}$ | 0,085 | 0,372 | 0,098 | 0,011 | 0,033 | 0,053 | 0,085 |
| $Fc^{Synchorotron}$ | 0,081 | 0,365 | 0,100 | 0,040 | 0,009 | 0,019 | 0,166 |
| **7RHO** | 0,064 | 0,410 | 0,099 | 0,001 | 0,100 | 0,005 | 0,251 |
| **Chain B** | | | | | | | |
| **SEM** | $P_{229}$ | $L_{231}$ | $E_{243}$ | $D_{250}$ | $Q_{280}$ | $N_{282}$ | $R_{286}$ |
| $Fc^{SFX}$ | 0,075 | 0,352 | 0,095 | 0,001 | 0,018 | 0,022 | 0,025 |
| $Fc^{Synchorotron}$ | 0,075 | 0,325 | 0,101 | 0,012 | 0,009 | 0,002 | 0,043 |
| **7RHO** | 0,087 | 0,400 | 0,115 | 0,013 | 0,128 | 0,017 | 0,241 |

### Description of Additional Supplementary Files

**Supplementary Movie 1:** Morph representation of the motion of Fc fragment of IgG1 subtype upon the binding of FcRn. To visualize the conformational changes, $Fc^{SFX}$ and $Fc^{FcRn}$ (PDB ID: 1FCC) structures are aligned and the movie is generated by using the Morph feature of *PyMOL.*

**Supplementary Movie 2:** Morph representation of the motion of critical residues in the FcRn binding site. To visualize the conformational changes, $Fc^{SFX}$ and $Fc^{FcRn}$ (PDB ID: 1FCC) structures are aligned and the movie is generated by using the Morph feature of *PyMOL.*

**Supplementary Movie 3:** Morph representation of the motion of Fc fragment of IgG1 subtype upon the binding of FcγR. To visualize the conformational changes, $Fc^{SFX}$ and $Fc^{Fc\gamma R}$ (PDB ID: 7URU) structures are aligned and the movie is generated by using the Morph feature of *PyMOL.*

**Supplementary Movie 4:** Morph representation of the motion of critical residues in the FcγR binding site. To visualize the conformational changes, $Fc^{SFX}$ and $Fc^{Fc\gamma R}$ (PDB ID: 7URU) structures are aligned and the movie is generated by using the Morph feature of *PyMOL.*

**Supplementary Movie 5:** Morph representation of the motion of Fc fragment of IgG1 subtype upon the binding of C1q. To visualize the conformational changes, $Fc^{SFX}$ and $Fc^{Fc\gamma R}$ (PDB ID: 6FCZ) structures are aligned and the movie is generated by using the Morph feature of *PyMOL.*

**Supplementary Movie 6:** Morph representation of the motion of critical residues in the C1q binding site. To visualize the conformational changes, $Fc^{SFX}$ and $Fc^{Fc\gamma R}$ (PDB ID: 6FCZ) structures are aligned and the movie is generated by using the Morph feature of *PyMOL.*

**Supplementary Movie 7:** Morph representation of the motion of critical residues in the glycan-binding site of Fc fragment of IgG1 subtype. To visualize the conformational changes during N-glycosylation, $Fc^{SFX}$ and $Fc^{Deglycosylated}$ structures are aligned and the movie is generated by using the Morph feature of *PyMOL.*

**Supplementary Movie 8:** Morph representation of the motion of critical residues in the glycan-binding site of Fc fragment of IgG1 subtype. To visualize the conformational changes during N-glycosylation, $Fc^{XRD(Ambient)}$ and $Fc^{Deglycosylated}$ structures are aligned and the movie is generated by using the Morph feature of *PyMOL.*

**Supplementary Movie 9:** Morph representation of the motion of critical residues in the glycan-binding site of Fc fragment of IgG1 subtype. To visualize the conformational changes during N-glycosylation, $Fc^{XRD(Cryogenic)}$ and $Fc^{Deglycosylated}$ structures are aligned and the movie is generated by using the Morph feature of *PyMOL.*

**Supplementary Movie 10:** Morph representation of the motion of critical residues in the glycan-binding site of Fc fragment of IgG1 subtype. To visualize the conformational changes during N-glycosylation, $Fc^{Synchrotron}$ and $Fc^{Deglycosylated}$ structures are aligned and the movie is generated by using the Morph feature of *PyMOL.*

**Supplementary Movie 11:** Time-lapse representation of 250 ns MD simulation for $Fc^{SFX}$ structure. The movie is generated by using the movie maker module of *VMD*.

**Supplementary Movie 12:** Time-lapse representation of 250 ns MD simulation for $Fc^{Synchrotron}$ structure. The movie is generated by using the movie maker module of *VMD*.

**Supplementary Movie 13:** Time-lapse representation of 250 ns MD simulation for $Fc^{Deglycosylated}$ structure. The movie is generated by using the movie maker module of *VMD*.
